# Navigating parasite antigen genetic diversity in the design of Plasmodium vivax serological exposure markers for malaria

**DOI:** 10.1101/2025.07.07.663616

**Authors:** Paolo Bareng, Kenneth W Wu, Eizo Takashima, Dionne Argyropoulos, Lauren Smith, Myo Naung, Ramin Mazhari, Kael Schoffer, Nicholas Kiernan-Walker, Anju Abraham, Macie Lamont, Somya Mehra, Pailene Lim, Jetsumon Sattabongkot, Wuelton Monteiro, Marcus Lacerda, Julie Healer, Chetan E Chitnis, Wai-Hong Tham, Takafumi Tsuboi, Ivo Mueller, Alyssa E. Barry, Rhea J Longley

## Abstract

**Background:** Plasmodium vivax poses a major obstacle to malaria elimination because it can lie dormant in the liver for weeks or months before reactivating and causing a relapse of infection. These dormant forms (hypnozoites) cannot be detected using standard diagnostics, but recent P. vivax exposure and by proxy, hypnozoite carriage, can be inferred using antibody-based tests (serological markers). In this study, we examined how genetic variation in P. vivax affects the utility of these antibody markers, and whether redesigned antigens could improve performance.

**Methods:** We analysed global P. vivax genetic data to assess variation in leading serological markers. Based on this, we produced new antigen versions (haplotypes) that better reflect global sequence diversity, compared to the commonly used reference strain (Sal-1). Antibody responses against these new constructs were then tested using samples from well-characterised cohorts in Brazil and Thailand. Antibody levels were assessed in relation to how recently participants had a qPCR-detectable blood-stage P. vivax infection. We compared the ability of the haplotypes and reference constructs to correctly identify individuals infected within the prior 9-months.

**Findings:** Extensive genetic diversity was identified in two P. vivax antigens, DBPII and MSP5. Several antigens had large numbers of circulating haplotypes globally, with the percentage with similar sequence identity to the reference Sal-1 ranging from 0.4% (MSP5) to 99% (S16). Two antigens exhibited strong differences in immunogenicity by region and construct (RBP2a and DBPII). However, for most proteins (5 out of 8), these differences had little impact on the accuracy of identifying recent exposure. In cases where performance was affected (e.g. RBP2a), this could be overcome by adding multiple antigens into the classification model.

**Interpretation:** Even highly diverse antigens can be effective serological exposure markers. Our findings highlight the importance of testing the impact of genetic diversity when designing serological tests and suggest practical strategies, such as using a mix of antigens, to ensure consistent performance across regions.

## INTRODUCTION

Over the last two decades, malaria incidence rates have substantially declined from 81 (in the year 2000) to 60 per 1000 population (in the year 2023)^1^. This impressive decline in malaria transmission is due to successful malaria control efforts, such as intensified vector control strategies and upscaled rapid diagnosis and treatment regimens, fueling optimism in several countries as they aim to eliminate malaria by 2030. However, Plasmodium vivax, one of the parasite species causing human malaria, is a hurdle to achieving this goal mainly because of its distinct biology, with the presence of an additional arrested stage in the liver (hypnozoites) that can cause relapsing malaria infection. Hypnozoites remain in the liver for weeks to many months following the initial mosquito bite induced infection and are major transmission reservoirs with up to 80% of all blood stage infections attributed to hypnozoite relapse^2,3^. Several pieces of evidence indicate that the proportion of P. vivax cases is increasing in areas where P. falciparum and P. vivax are co-endemic^4,5^, signifying that more robust control and elimination strategies, including surveillance, are needed to reach the ambitious malaria elimination goals.

Several countries have employed active interventions alongside other control strategies, to alleviate the burden of P. vivax transmission. An example is mass drug administration (MDA), which entails treating all individuals, whether infected or not, to clear current infections and eliminate potential hypnozoites in the liver^6^. Significant drawbacks exist, such as the safety, efficacy and community acceptability of the antimalarials and the potential to accelerate drug resistance^7,8^. For P. vivax, safety is a particular concern for MDA. Mutations in the enzyme G6PD in human populations are protective against malaria^9^, however, they render those individuals more susceptible to hemolysis upon treatment with 8-aminoquinolines, the current frontline treatment for P. vivax liver-stages. An alternative strategy, mass screening and treatment (MSAT) has been proposed where individuals undergo screening for active blood-stage P. vivax infections before treatment with antimalarial drugs. Although MSAT may have certain advantages compared to MDA in terms of reducing overtreatment and the effects of drug resistance, the success of this strategy still depends on the population coverage and the tool/s used in screening infections. Current diagnostic tools such as microscopy and rapid diagnostic tests (RDTs) are not sensitive enough to detect the low-parasite density infections common with P. vivax nor detect hypnozoites^10,11^. Alternatively, polymerase chain reaction (PCR) is more sensitive but also cannot detect hypnozoites in the liver. Therefore, this gap in P. vivax diagnostics highlights the need for more efficient tools capable of detecting individuals carrying hypnozoites, for both surveillance and the implementation of targeted interventions to rapidly reduce P. vivax transmission.

A previously published study used total IgG antibody levels against a panel of serological exposure markers (SEM) to identify individuals with recent P. vivax infection^12^. The principle underlying this assay is that the measurement of antigen-specific IgG antibodies can inform on past and present exposure to infection^13^. Based on hypnozoite relapse biology, nearly all primary relapse infections occur within the first 6-9 months following the initial mosquito bite induced infection^14^. Thus, SEMs that can inform on P. vivax exposure within the prior 9-months can potentially classify individuals with hypnozoite carriage. This study developed a rigorous analytic pipeline to evaluate over 300 P. vivax antigens, resulting in the final panel of 8 candidate proteins with sensitivity and specificity of 80%^12^. This SEM assay can be used for surveillance and for serological testing and treatment (seroTAT)^15^, specifically targeting individuals for P. vivax radical cure^16,17^. Most importantly, seroTAT could help prevent the blanket approach of MDA, ensuring a more efficient and targeted intervention.

Despite the promise of SEMs as a potential public health intervention, the effects of genetic diversity on the utility of these diagnostic biomarkers have not been addressed. The proteins expressed to measure antibody responses in the development of these sero-diagnostic tools were based on a single P. vivax reference strain (Sal-1), representing only a small proportion of parasites in natural parasite populations^18–20^. Therefore, despite overall good classification performance of the SEMs across geographic regions^12^, the observed differences in antibody levels among the analysed cohorts (Thailand, Solomon Island, and Brazil) could be due to strain-specific responses elicited by the Sal-1-based antigen panel. Analysing the role of parasite genetic diversity in generating strain specific responses is crucial for the development of highly sensitive sero-surveillance tools that work equally well across different geographic regions with varying parasite populations. Whilst progress has been made assessing the impact of antigen genetic diversity on Plasmodium vaccine-induced immune responses ^21^, this has not been tested in the context of serological exposure markers where only antibody magnitude (and not function) is important.

Here, patterns of genetic diversity and immune selection pressure of the leading SEM antigens, including region II of Duffy binding protein (DBPII), merozoite surface protein 5 (MSP5), merozoite surface protein 1 (MSP1_19_), merozoite surface protein 8 (MSP8), reticulocyte binding protein 2a (RBP2a), reticulocyte binding protein (RBP2b), RH5-interacting protein (RiPR), rhoptry-associated membrane antigen (RAMA), Plasmodium translocon of exported protein 150 (PTEX150), tryptophan-rich protein (Pv-Fam-a), and sporozoite traversal protein (S16) were investigated. Sequence diversity was calculated for both linear sequence and three-dimensional protein structure using global whole genome sequence data obtained from P. vivax sequencing projects carried out by the Malaria Genetic Epidemiology Network (MalariaGEN) and Broad Institute^22,23^. The population genetic data was then used to strategically select antigen haplotypes predicted to cover most of the antigenic diversity observed, with these variant constructs expressed and tested for immunological reactivity and classification performance compared to the Sal-1 reference strain.

## METHODS

### Selection of P. vivax SEM antigens

An initial panel of 14 P. vivax SEM antigens were selected based on prior work, including the top 8 antigens used in the published algorithm^12^ along with other high performing candidates^24^. Three antigens (PVX_097720 – MSP3a, KMZ83376.1 – EBP, PVX_112670 – a Pv-fam-a) fell into the blacklisted regions of the PvP01 genome (part of the genome near telomere and sub-telomere regions)^25^ and are either part of the hypervariable region or not part of the nuclear chromosome (bed file genomic coordinate available at https://github.com/paolobareng/pvp01_blacklisted_regions.git). The remaining 11 P. vivax antigens were taken through the below pipeline (Figure S1). When specific regions of these proteins were assessed, this was based on the amino acid sequences used in the original Sal-1 constructs^12^ (mapped to the PvP01 genome), and these are provided in Table S1.

Whole genome sequences (WGS) and variant calling The meta-population sequence data was used to derive SEM antigen sequences for the 11 P. vivax antigens. Pre-processing of the WGS data^26^ (n=613 genomes) from MalariaGEN P. vivax Genome Variation Project 2016, sequenced using Illumina short-read technology at the Sanger Institute and the Broad Institute (USA), was carried out ^22,23^ using a bioinformatics pipeline according to the GATK (version 4.0.12.0) best practices workflow^26^. The sequences were mapped against P. vivax P01 reference genome (PlasmoDB release 50: 2020/12/15). The extracted WGS samples are available at: https://github.com/paolobareng/Pvivax_SEM. To retain variants within the 14 nuclear chromosomes, we removed variants on shorter contigs (n=226), mitochondrial, and apicoplast genomes. Then, we filtered out single nucleotide polymorphisms (SNP) located within the subtelomeric hypervariable, subtelomeric repeats, centromeric regions, and indels within the nuclear chromosomes, as these variants have different evolutionary mechanisms that are not of interest to our study. Afterwards, we performed another variant and sample filtration using stricter quality metrics. Variants and samples were retained if they met the following criteria: Missingness rate (by sample) > 0.03; Quality by depth (QD) > 20; Mapping quality (MQ) > 50; Mapping quality rank suum test (MQRankSum) >-2; Symmetric odds ration (SOR) < 1;-4 < ReadPosRankSumTest < 4; Missingness rate (by site) > 0.1; and countries with at least n=5 samples. Moreover, F_WS_ metric (utilising moimix R package^27^) was used to infer multiplicity of infection (MOI). WGS were classified as clonal (MOI = 1, FWS > 0.95), multiclonal (MOI = 2, 0.80 < FWS ≤ 0.95), or of indeterminate clonality (MOI > 2, FWS ≤ 0.80). Only major clones were included in the final dataset, with a minimum coverage depth of 5x for MOI=1 and 10x for MOI=2 infections.

Genes of interests were extracted from the final WGS data using the following annotations: PVP01_0623800 (DBP), PVP01_1402400 (RBP2a), PVP01_0800700 (RBP2b), PVP01_0816800 (RiPR), PVP01_0418400 (MSP5), PVP01_1032600 (MSP8), PVP01_0107500 (RAMA), PVP01_1313200 (PTEX150), PVP01_0728900 (MSP1_19_), PVP01_0202200 (Pv-fam-a), PVP01_0305600 (S16). The resulting variant call format (VCF) was then converted to Fasta file format for downstream population genetic analyses (see further details at https://github.com/myonuang/Naung-et-al-2021). Further data cleaning, such as removal of singletons or sequencing artifacts, was performed to improve its overall quality.

### Antigen diversity analyses

Traditional measures of population genetics and Tajima’s D test of neutrality were calculated using a custom R-package VaxPack (https://github.com/BarryLab01/vaxpack). Measures included: the nucleotide diversity (π), which is the average number of allele differences per site between any two random sequences, haplotype diversity (Hd), which is the probability that the two randomly chosen haplotypes are different, the number of polymorphic or segregating sites, number of synonymous SNPs (SP), and non-synonymous SNPs (NS). Country-level populations with more than 15 sequences were calculated for deviations from neutrality using the Tajima’s D statistic. A negative Tajima’s D value signals an excess of rare alleles suggestive of purifying selection or recent population expansion, while a positive Tajima’s D indicates intermediate allele frequency suggestive of balancing selection, possibly linked to host immune pressure. To visualize the genetic relatedness of sequence data in different parts of the world, haplotype network diagrams were constructed for non-synonymous polymorphisms in each antigen gene using PoPArt software (v. 4.8.5)^28^. Diversity was assessed only within the region of the protein used in the P. vivax SEM tool.

The AlphaFold algorithm^29^ was used to model three-dimensional (3D) protein structures of PvP01 DBPII, RBP2a, and RBP2b. To assess the accuracy of the predicted models, we overlaid the Alphafold structures with the corresponding experimentally defined crystal structures available for the Sal-1 strains (Figure S6D). Then, a modified Biostructmap Python script was used to calculate the population genetic measures and Tajima’s D over the predicted protein structures in a 15 Å radius sliding window. Visualisation of the structural genetic diversity and selection pressure was done with PyMOL software^30^.

### Selection and expression of P. vivax antigen haplotypes

For the 11 P. vivax antigens taken through the bioinformatics pipeline, we selected 1-3 haplotypes per antigen based on haplotype network maps and the presence of global isolates covered as detailed in the Results. A total of 14 haplotypes were selected for 8 unique P. vivax SEMs, these were expressed, along with use of the Sal-1 constructs as the reference (already expressed as previously described^12^). For three P. vivax SEMs (MSP1_19_, RAMA, S16), the selected haplotype had no amino acid differences compared to Sal-1 and thus was not expressed nor tested further immunologically (Figure S1). The selected haplotypes were expressed using the wheat-germ cell-free (WGCF) protein expression system using previously described standard methods^31^, and protein was purified with a C-terminal His-tag by single-step affinity purification^12^. Purified recombinant protein was applied to sodium dodecyl sulphate polyacrylamide gel electrophoresis (SDS-PAGE) and visualised with Coomassie Brilliant Blue (CBB) staining (Figure S2). The final list of antigens tested in the immunological experiments and their notation is listed in Table S2.

### Total IgG antibody assay

All selected antigens (Table S2) were coupled to BioRad COOH magnetic beads as previously described^32^. Briefly, the carboxylic acid functional groups on the magnetic microspheres were activated using sulfo-NHS and EDC before incubating with the chosen proteins in PBS overnight. The protein amount was optimized against a standard curve created with a positive control pool from Papua New Guinea (PNG), and is provided in Table S2.

Total IgG was measured in all samples using a multiplexed Luminex-based assay, as previously described^32^. Briefly, diluted plasma samples (1:100) were incubated with 0.1 ml of each coupled microsphere on a plate shaker for 30 minutes. IgG binding to coupled proteins was detected with goat anti-IgG PE-labelled secondary antibody (1:100 dilution) after incubation for 15 minutes on a plate shaker. Plates were read on a MAGPIX Luminex instrument. The median fluorescent intensity (MFI) output was converted to relative antibody units (RAU) using the positive control PNG pool standard curve. A 2-fold 10-point serial dilution from 1/50 to 1/25,600 enabled fitting of a five-parameter logistic regression standard curve as previously described^32^.

### Ethical approvals

The Human Research Ethics Committee at WEHI (Australia) approved the use of collected samples in Melbourne (14/02), and approved collection and/or use of the negative control samples (14/02). The Ethics Committee of the Faculty of Tropical Medicine, Mahidol University (Thailand) approved the 12-month observational Thai cohort (MUTM 2013-027-01). The Brazilian National Committee of Ethics (Brazil) and the Ethics Committee of the Hospital Clinic, Barcelona (Spain) approved the 12-month Brazilian cohort (349.211/2013 and 2012/7306). All participants gave informed consent and/or assent to participate in the study.

### Human study samples: negative controls and observational cohorts

Plasma samples previously collected from malaria-naïve individuals were used to determine the background antibody response not induced by infection, and to act as negative controls in the classification model. A total of 370 samples from the Australian Red Cross (ARC) (n=100), the Brazil Red Cross (BRC) (n=96), the Volunteer Biospecimen Donor Registry (VBDR) at WEHI (n=102), and the Thai Red Cross (TRC) (n=72) were used, as previously described^12^. ARC and VBDR sample sets were both from malaria-free Melbourne, Australia. BRC samples were collected from the Rio de Janeiro State Blood Bank, a non-endemic region with no local malaria transmission since the 1960s. The TRC excluded individuals from donating blood if they had a confirmed malaria infection in the last 3 years or had been in malaria-endemic regions a year prior.

Plasma samples previously collected from two year-long observational cohort studies in Thailand^33^ and Brazil^34^ were used as our P. vivax exposed/endemic samples. We utilized samples from the last visits of these cohorts (n=547-774 Thailand – dependent on antigen due to low plasma availability, n=923 Brazil), as previously described. The age range of participants was 0–103 years in the Brazil cohort and 1–78 years in the Thailand cohort. The self-reported gender distribution in the Brazil cohort was 50.6% women and 49.4% men, while the Thailand cohort was 54.8% women and 45.2% men participants. After enrolment, individuals were sampled monthly with active detection of Plasmodium blood-stage infection by PCR. Using this PCR data, individuals were classified as currently infected at the time of antibody measurements, infected within the prior 9-months, infected 9-12 months ago or no detected infections throughout the yearlong cohort period.

### Statistical analysis: classification of recent exposure

Individuals in the malaria-endemic cohorts were defined as either recently infected with P. vivax in the prior 9-months or not, utilizing the monthly PCR data. Individuals from the malaria-naïve samples were classified as not exposed. The ability of total IgG antibodies to classify individuals as recently exposed to P. vivax in the prior 9-months was assessed using binary cut-offs with the results depicted in Receiver-Operating-Characteristic (ROC) curves. To assess the significance of differences, the statistical comparison of the Area Under the Curve (AUC) between the reference strain antigens and their genetic haplotypes was performed using bootstrap resampling (2000 replicates), with each replicate generated by sampling with replacement from the original dataset, maintaining the original sample size. The analysis was conducted in R using the pROC package^35^.

To explore the top combinations of P. vivax antigens, a random forest algorithm was created for all possible combinations using a 10-fold cross validation with five repeats. The random forest was fit using the ranger v0.17.0 R package^36^ with 1,000 trees and all other hyperparameters were default. The top-performing combinations of the two to eight antigens were selected and classification performance was evaluated using the AUC. This analysis was performed using the RBP2a Sal-1, SEM 28 and SEM 29 haplotypes for the combined dataset including the Thai and Brazilian cohorts and negative control samples, as well as for the Thai and Brazilian datasets separately. Multi-antigen combinations greater than eight were not assessed due to the minimal gains in classification performance for the overall dataset, also observed in Longley et al., (2020).

## RESULTS

Global sequence data revealed extensive diversity in genes encoding DBPII and MSP5 MSP5 (πx10^-3^=14.8) and DBPII (πx10^-3^=7.7) have the highest global diversity among the P. vivax SEM proteins (Table S1). The high genetic diversity of the genes encoding these proteins was consistent in all analysed populations (Figure 1, Table S1). By contrast, other antigens localised on the merozoite secretory organelles (RBP2a, RBP2b, RAMA, and RiPR), the merozoite surface (MSP8), as well as other newly discovered proteins (PTEX150 and Pv-fam-a), demonstrated low nucleotide diversity (Figure 1, Table S1). Similarly, MSP1_19_ and the gametocyte antigen S16 exhibited low genetic diversity, with only 1 to 2 polymorphic sites in some populations. Even despite having relatively few polymorphisms, several antigens such as RBP2a, RBP2b, RiPR, and PTEX150, have relatively high haplotype diversity, indicating that rare alleles are generated during genetic recombination events, leading to a high number of novel haplotypes circulating in the population (Figure 1).

**Figure 1.**
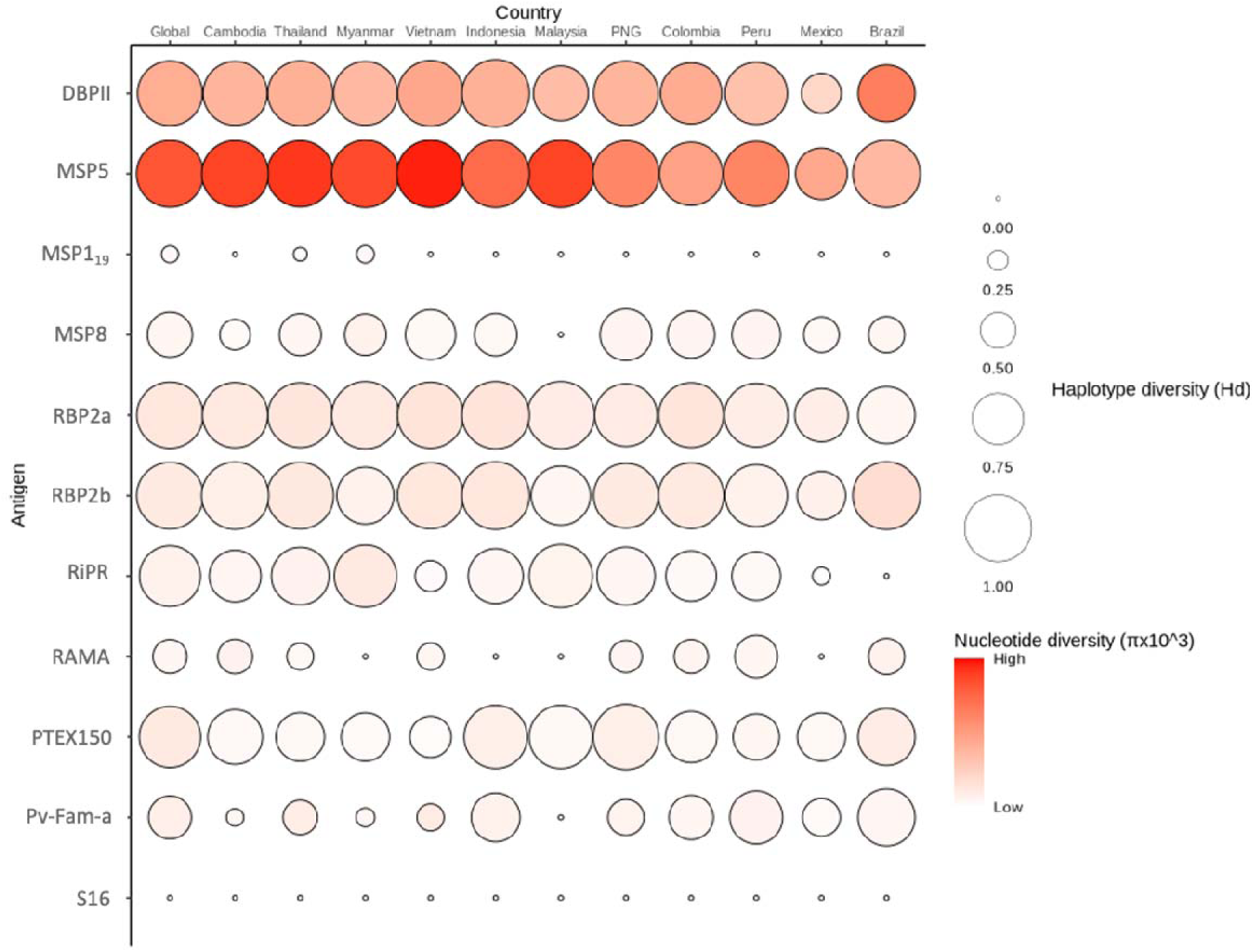
**Nucleotide (**π**) and haplotype diversity (Hd) in the expressed regions of *P. vivax* SEM antigens across diverse populations.** The darker color gradient indicates higher π, and the circle’s size corresponds to Hd, with larger nodes representing higher Hd. Diversity was only assessed within the expressed region of the protein used in the P. vivax SEM tool.

The level of nucleotide diversity across the length of the genes found amongst all sequences globally was calculated (Figure 2). The major pattern observed was peaks of diversity in specific domains or regions of each gene, and this was consistent across all populations analysed (Figure S3), suggesting that immune selection could be responsible for maintaining this domain-specific variability. For example, region II in the gene encoding DBPII exhibits greater polymorphism than subdomains I and III, and this pattern is observed in all populations. Additionally, high nucleotide diversity was observed in exon 1 of MSP5, whilst exon 2 was generally conserved. Small peaks of nucleotide diversity were also detected in specific regions of other SEM antigens, including the N-terminal of the genes encoding RBP2a, RBP2b, RiPR, and Pv-fam-a, as well as both the ASN-rich region and EGF domain of MSP8. Furthermore, for three antigens where crystal structures were available in the Protein Data Bank (RBP2a:4Z8N; RPB2b:5W53; DBPII:4Y5S), thus enabling verification of AlphaFold predicted structures, “spatial” diversity was calculated revealing areas of high nucleotide diversity within specific parts of the structures (Figure S4A).

**Figure 2.**
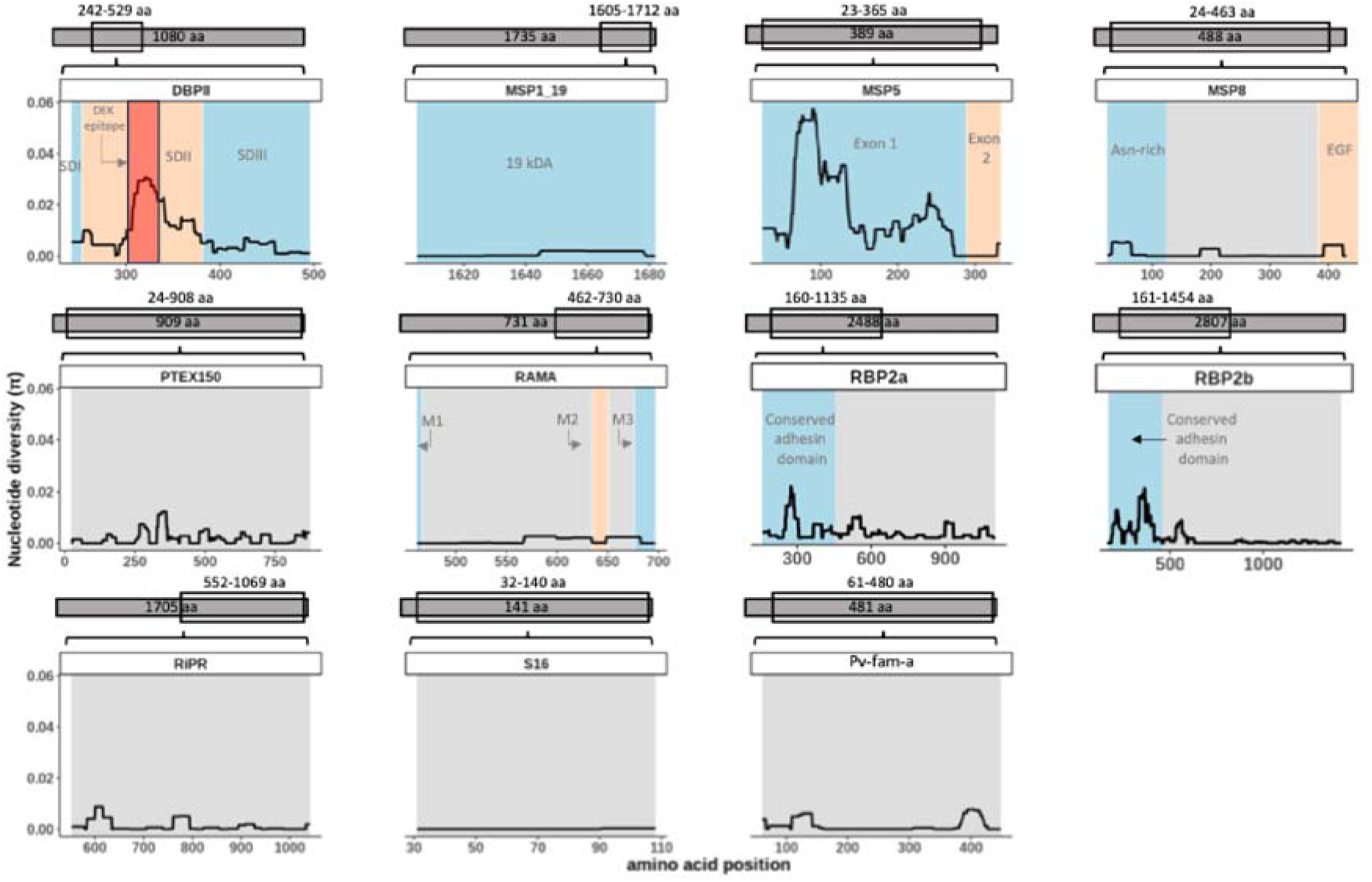
Global nucleotide diversity along the length of all 11 *P. vivax* genes. Nucleotide diversity across the gene sequence. Upper schematic highlights the region of the gene included in the expressed P. vivax protein constructs. Coloured blocks highlight different domains based on previous studies, with grey indicating undefined regions. Nucleotide diversity (π) was calculated in sliding window approach (window size=100; step size=3).

### Polymorphic gene domains exhibited evidence of positive balancing selection

Comparative analysis of Tajima’s D index revealed predominantly positive values across most populations for the highly variable DBPII and for MSP5 (Table S1). This implies that balancing selection pressure that maintains diversity among these populations. The exception was for Mexico which had negative Tajima’s D values for DBPII, and PNG for MSP5, indicating directional or purifying selection. In contrast, Tajima’s D values in the remaining SEM antigens (MSP1_19_, MSP8, RBP2a, RBP2b, RiPR, RAMA, PTEX150, Pv-fam-a, S16) were either negative or had insignificant departures from neutrality, which is consistent with their low genetic diversity. Additionally, there was no discernible differences in Tajima’s D patterns in these low-diversity SEM antigens in different geographical regions, suggesting similar selective pressures are acting across populations. When assessing Tajima’s D statistics across the entire constructs (Figure S5), positive peaks were identified in the polymorphic region II of DBPII and the genetically diverse N-terminal of MSP5. Despite differences in the selection pressures acting in various regions of DBPII and MSP5, the highly variable regions exhibit positive Tajima’s D, suggesting that immune selection could be a major driving force for allelic diversity in these domains. Significant signatures of balancing selection were detected in the known conserved adhesion domain of RBP2a and RBP2b, in certain populations (Figure S5). Conversely, antigens such as MSP1_19_, MSP8, RiPR, RAMA, PTEX150, Pv-fam-a, and S16 mostly displayed either neutral or negative selection, with occasional peaks of balancing selection in some regions (Figure S5). This pattern aligns with the expected selection pressure, considering the low genetic variability observed in these antigens.

Tajima’s D estimates were incorporated into the modelled protein structures of DBPII, RBP2a, and RBP2b (Figure S4B), to investigate and compare structural patterns of selection pressure across various populations. DBPII showed strong signals of balancing selection particularly within the inhibitory region called the “DEK” epitope (Figure S6A); this epitope is known to be highly variable and lies directly opposite Duffy antigen (DARC)^37^. While parasite populations in Cambodia, Thailand, Colombia, and Peru displayed significant Tajima’s D around the DEK epitope (Figure S4B), sequences from PNG and Mexico showed low signals of Tajima’s D, indicating different types of selection pressure acting upon this subdomain. RBP2a displayed positive Tajima’s D in the conserved adhesin region with high rates of allelic polymorphism (encircled in Figure S4B), except for Mexico where weak positive signals of Tajima’s D were detected. The variable region of the RBP2b, close to the TfR1 binding sites (Figure S6C), appeared to have low to negative Tajima’s D value in most populations (Figure S4B), suggesting evidence of neutral or directional selection. Nevertheless, the parasite population in Mexico stood out, as the structural patterns of selection pressure greatly differed from those of other countries (Figure S4B).

### Haplotype networks and design of new constructs for immunological testing

To investigate the genetic relatedness of the antigen sequences, haplotype networks (Figure 3) were constructed using non-synonymous polymorphisms from specific gene constructs (i.e. the region of the protein included in the P. vivax SEMs). A complex network of haplotypes was expected for the genetically diverse antigens DBPII and MSP5, as higher rates of polymorphism generate a higher number of haplotypes in the population. Meanwhile, RBP2a and RBP2b network analyses lack population structure despite having low nucleotide diversity. These haplotype networks confirm the high haplotype diversity values computed in these genes (Table S1). PTEX150 showed some geographical clustering, with sequences belonging to specific regions of Asia, America, and PNG tending to group. MSP1_19_, MSP8, RAMA, Pv-fam-a, RiPR, and S16 exhibited less structured haplotype networks, consistent with their observed low genetic variation.

**Figure 3.**
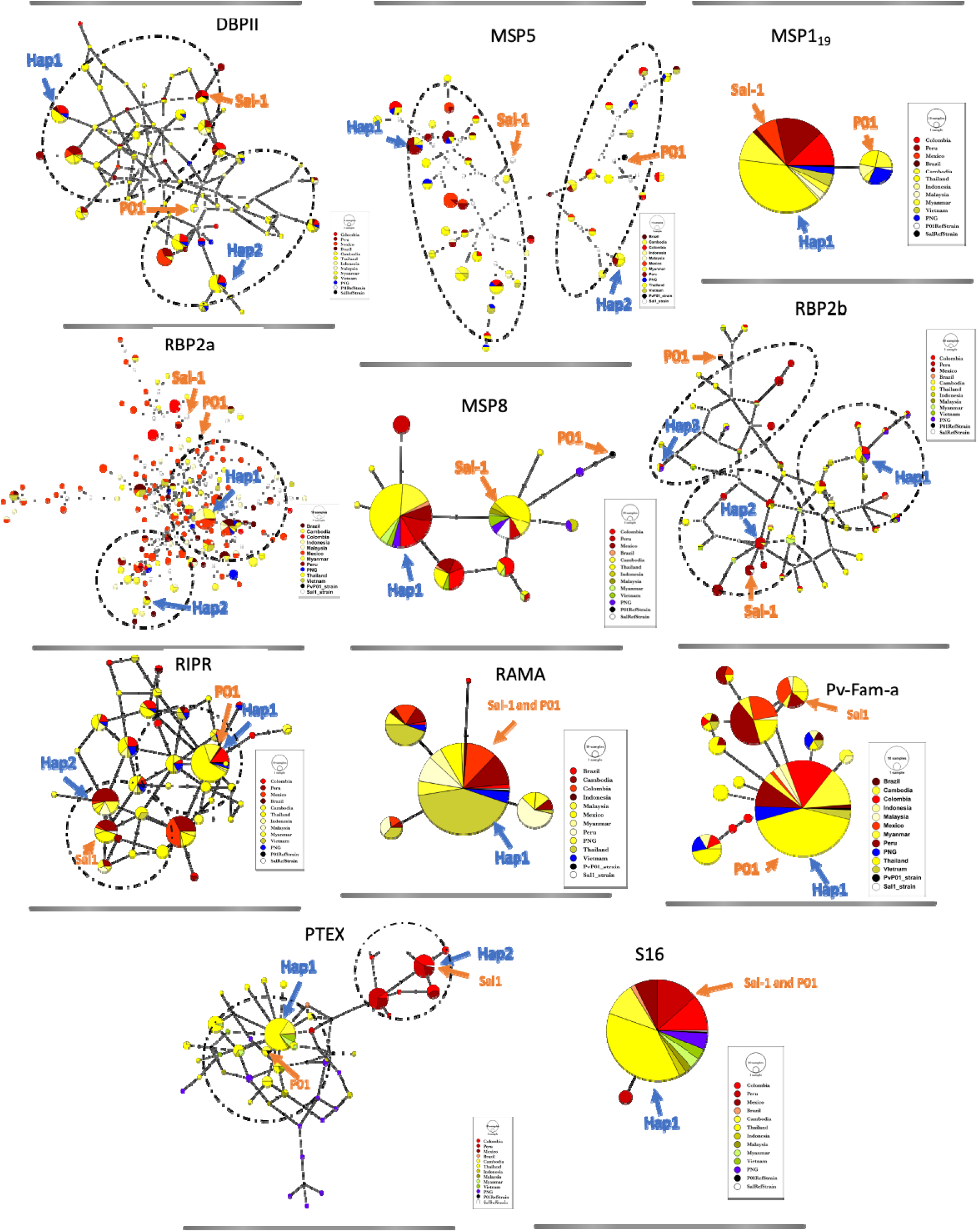
Haplotype network plots and the distribution of diversity-covering haplotypes for each antigen. Each node in the plots represents a unique haplotype, with the node size corresponding to haplotype frequency, and the color indicating specific country (refer to the legend). The red arrow denotes the known reference alleles Sal-1 and P01, whereas the blue arrows indicate selected haplotypes that could cover the diversity observed in these antigens. Dotted circles represent identified clusters for highly diverse antigens.

The Sal-1 reference, which was the strain used in designing P. vivax SEMs, was either in very low frequencies (<4%) for genes with high haplotype diversity (DBPII, MSP5, RBP2a, RBP2b) or grouped within a geographical cluster, as observed for RiPR (7%) and PTEX150 (10%) (Table 1, Figure 3). Higher proportions of haplotypes were similar to Sal-1 for the genes encoding Pv-fam-a, MSP8, RAMA, MSP1-C terminal, and Pvs16 (14-99%). Drawing on the diversity measures outlined in this study, the following criteria for the selection of haplotypes were established to maximise diversity coverage: 1. The highest frequency haplotype/s in the population, 2. The highest frequency haplotype in the identified clusters. Figure 3 shows the location of the selected sequences in the haplotype network diagram (blue arrow), while Table 2 summarises the population frequency of diversity-covering haplotype/s for each antigen and their mutational difference to Sal-1. Of these, the haplotypes selected for MSP1_19_, RAMA, and S16 had no mutation differences compared to Sal-1 reference and thus were not expressed. For the same reason, Hap2 for PTEX150 was not expressed. The final panel of antigens tested immunologically are listed in Table S2.

**Table 1.** Proportion of identified haplotypes in the population similar to the Sal-1 reference strain used in the P. vivax SEMs. Antigens listed in order of lowest-highest similarity.

| Antigen (amino acid position) | % of Haplotypes similar to Sal-1 |
| --- | --- |
| MSP5 (23-365) | 0.4 |
| RBP2a (160-1135) | 0.4 |
| RBP2b (161-1454) | 2 |
| DBPII (242-529) | 4 |
| RiPR (552-1075) | 7 |
| PTEX150 (24-908) | 10 |
| Pv-fam-a (61-280) | 14 |
| MSP8 (24-463) | 22 |
| RAMA (462-730) | 71 |
| MSP1<sub>19</sub> (1605-1712) | 89 |
| S16 (31-140) | 99 |

**Table 2:** List of selected haplotypes derived from the genetic diversity analysis. Information includes the number of mutation differences from Sal-1 reference allele, the frequency of the sequence in the population, and the total proportion of diversity covered in the population. The selected sequences are available at the following link: https://github.com/paolobareng/Pvivax_SEM.

| Antigen | Haplotype | Construct (aa) | no of mutation difference/s to Sal-1 | Population frequency (%) | Total population covered (%) |
| --- | --- | --- | --- | --- | --- |
| DBPII | Hap1 | 242-529 | 4 | 8.3 | 16.1 |
|  | Hap2 |  | 11 | 7.8 |  |
| MSP5 | Hap1 | 23-365 | 29 | 4.3 | 6.8 |
|  | Hap2 |  | 31 | 2.5 |  |
| MSP1<sub>19</sub> | Hap1 | 1622-1729 | 0 | 89.5 | 89.5 |
| RBP2a | Hap1 | 160-1135 | 7 | 9.3 | 11.1 |
|  | Hap2 |  | 8 | 1.8 |  |
| RBP2b | Hap1 | 161-1454 | 8 | 8.2 | 13.4 |
|  | Hap2 |  | 1 | 3.7 |  |
|  | Hap3 |  | 13 | 1.5 |  |
| MSP8 | Hap1 | 24-463 | 1 | 54.5 | 54.5 |
| RiPR | Hap1 | 552-1075 | 3 | 23.6 | 34.4 |
|  | Hap2 |  | 1 | 10.7 |  |
| RAMA | Hap1 | 462.73 | 0 | 70.8 | 70.8 |
| Pv-fam-a | Hap1 | 61-421 | 9 | 60.2 | 60.2 |
| PTEx | Hap1 | 24-908 | 5 | 25.5 | 37.5 |
|  | Hap2 |  | 0 | 12 |  |
| S16 | Hap1 | 31-110 | 0 | 98.5 | 98.5 |

### Immunoreactivity of selected haplotypes versus Sal-1 reference in Thai and Brazilian human cohorts

IgG antibody responses were measured against all selected haplotypes and the reference Sal-1 strain at the end of year-long observational cohort studies conducted in Brazil and Thailand. We compared immunogenicity against the haplotypes compared to the reference (Figure 4), expecting higher IgG signal in individuals with current P. vivax infections, followed by those infected 1-9 months ago then 9-12 months ago, and with lower signal in those with no detected Plasmodium infections over the yearlong period and in malaria-naïve controls. We also paid attention to background in the malaria-naïve controls, as this could signify protein quality^12^ rather than an inherent cross-reactive response to the haplotype. Similar patterns were observed between countries and between the reference sequence and haplotypes for MSP5, RiPR, PTEX150, Pv-fam-a and RBP2b. High background was evident against the MSP8 haplotype in the controls, despite the protein appearing high purity (Figure S2B). Two proteins exhibited strong differences in immunogenicity by both country and reference compared to haplotype: RBP2a and DBPII. In Thailand, the strongest peak immunogenicity and expected step-wise decline in IgG level with increasing time since last detected P. vivax infection was observed against RBP2a Hap1, however against the Sal-1 reference strain for Brazil. Conversely, all DBPII constructs (Sal-1, AH, and two haplotypes), were strongly immunogenic in Brazil, however none gave strong expected patterns in Thailand.

**Figure 4.**
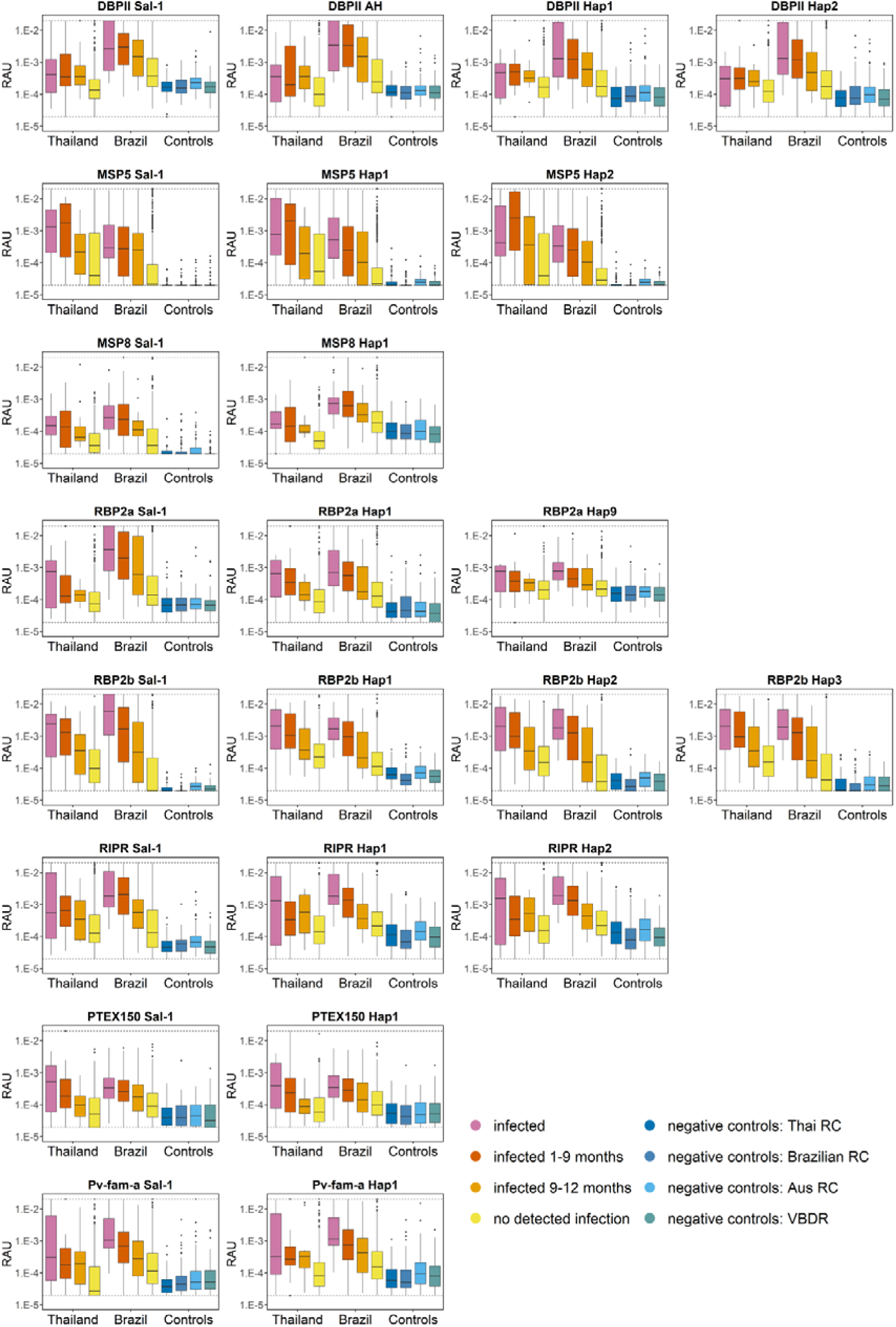
IgG antibody levels in Thai and Brazil individuals against the *P. vivax* serological exposure marker proteins comparing the reference strain (Sal-1) and the identified haplotypes. IgG levels were measured at the final visit of yearlong cohorts in Thailand and Brazil. Monthly bleeds enabled qPCR-detection of Plasmodium infections and calculation of the time since last detected P. vivax blood-stage infection. Participants were stratified by this calculation: current P. vivax infection (n=13-20 for Thai and n=39 for Brazil), P. vivax infection 1-9 months ago (n=21-36 for Thai and n=165 for Brazil), P. vivax infection 9-12 months ago (n=12-19 for Thai and n=31 for Brazil), no detected P. vivax infections (n=501-699 for Thai and n=688 for Brazil). IgG levels were also measured in four panels of negative controls, Australian Red Cross (n=100), Volunteer Biospecimen Donor Registry (n=102), Brazil Red Cross (n=96) and Thai Red Cross (n=72).

In each cohort, IgG levels were generally positively correlated amongst the different constructs of the one SEM antigen (Figure S7). The exception was in Thailand for DBPII and RBP2a (Figure S7B), with spearman r values less than 0.7. There were also relatively strong correlations between various RBP2b and MSP5 constructs in both cohorts, signifying potential co-acquisition of these antibodies.

### Classification performance of selected haplotypes versus Sal-1 reference in Thai and Brazilian human cohorts

Measured IgG antibody responses to each protein individually were then utilized to classify recent exposure to P. vivax infections in the prior 9-months, with the monthly qPCR data used as the reference or truth. Classification performance was assessed as sensitivity and specificity with the results presented in receiver operator characteristic curves (Figure S8), and then comparison of the classification when using the reference strain or the selected haplotypes by comparing the area under the curve summary statistic (Figure 5, Table S3). For most P. vivax proteins (5/8) there was no significant impact on classification performance when using the Sal-1 reference strain or one of the haplotype variants. For two P. vivax SEM proteins, MSP8 and RBP2a, there were significant differences in both Thailand and Brazil. For MSP8 the reference Sal-1 antigen had higher classification performance in both cohorts. However, for RBP2a, Hap1 had the highest AUC in Thailand whereas Sal-1 had the highest AUC in Brazil. The haplotype network analysis did not reveal significant geographical clustering for RBP2a (Figure 3). In Brazil only, there was slightly better performance for RBP2b when using Hap3 compared to Sal-1, however even Sal-1 had an AUC >0.8, likely limiting any impact on effectiveness^38^.

**Figure 5.**
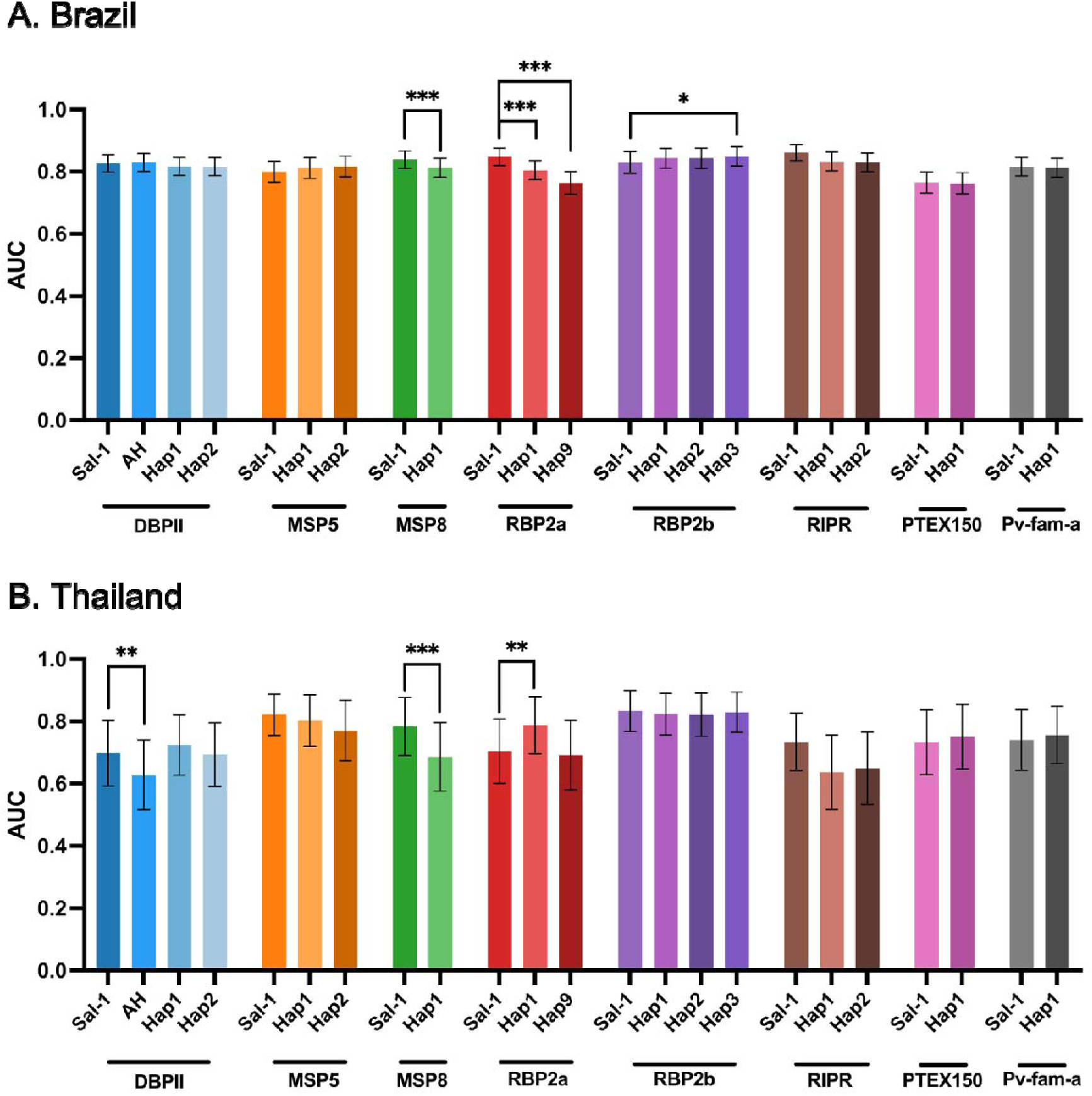
Summary performance for correctly classifying recent exposure to *P. vivax* when using either the reference or variant haplotypes selected. Results are presented as area under the curve (AUC) of the ROC curves presented in Figure S8, with 95% confidence intervals. Statistical difference between classification for each antigen when using the reference (Sal-1) or the identified haplotype variants is shown. Statistical significance was assessed using bootstrap resampling (2000 replicates). Results were considered significant at p < 0.05 (*), p < 0.01 (**), and p < 0.001 (***).

In the final test, antibody responses to the 8 P. vivax SEM antigens are used in combination and not individually^12^, as this increases classification performance. Using RBP2a as an example, we tested combinations of two to 8 antigens to test the impact of genetic diversity on a single antigen once it is used in combination. When RBP2a Sal-1 or either haplotype were combined with another antigen (Sal-1 reference strain), differences in AUC were still observed, however once a third antigen was added the detrimental impact of using either haplotype was diminished (Figure 6A). This pattern was also observed when Thai and Brazilian datasets were analysed separately (Figure 6B, C), however in Thailand the AUC declined after more than five antigens were included (Figure 6C), likely due to model overfitting caused by the limited number of seropositive samples available to train the model. The different antigens utilized in these combinations are available in Table S4.

**Figure 6.**
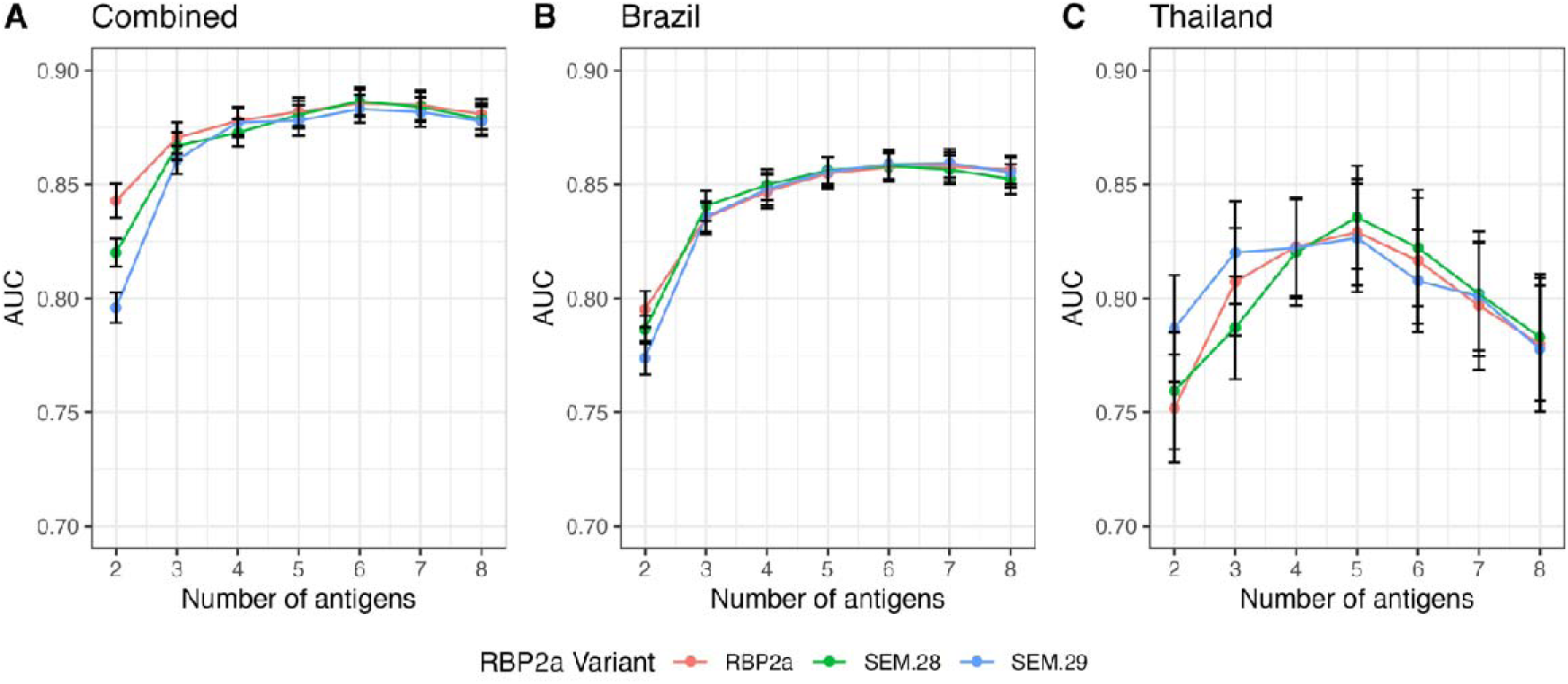
Impact of RBP2a genetic diversity when used in combination with other *P. vivax* SEMs. Performance of the top eight P. vivax antigens in classifying seropositive infection within the last nine months was measured by the cross-validated AUC of the ROC curve values generated from the random forest classifier. Results are shown for the instances where the model was trained and tested on (A) the combined dataset, including cohort studies from Brazil and Thailand along with negative control samples, and separately on (C) the Brazilian cohort and (C) the Thai cohort. The analysis was conducted using the RBP2a Sal-1 (pink), SEM 28 (green) and SEM 29 (blue) haplotypes.

## DISCUSSION

Developing a sero-surveillance assay necessitates a comprehensive evaluation of antigens, as antibody response profiles, longevity, and sequence diversity can differ among them. We previously proposed the “seroTAT” approach, a test and treat strategy employing a panel of P. vivax serological exposure markers to identify individuals with hypnozoites^12^. This approach represents the first method to indirectly identify hypnozoites carriers and has led to the first WHO Preferred Product Characteristics for a test that can detect relapse risk^39^, paving the way for new public health approaches to accelerate malaria elimination. Whilst the P. vivax SEMs were developed and validated in three discrete geographic regions (Thailand, Brazil, Solomon Islands)^12^, and have since been applied in Peru^40^ and Cambodia^41^, IgG antibody levels still vary in different populations and this could be due to various factors including antigen diversity. Therefore, genetic diversity data should be factored into the development of any antigen-based tool, as the inclusion of highly diverse genes can potentially compromise the sensitivity of the assay.

The present study addresses this concern by examining the linear (primary sequence alignment) and structural patterns of genetic diversity and selection pressure of the SEM antigens. Furthermore, this study then tests the utility of different antigen variants that are more representative of global parasite populations. Ultimately, this led to one of the original top eight antigens to be dropped from the sero-surveillance tool; MSP3a, due to its large size and complexity (large number of haplotypes and high nucleotide diversity). Of the remaining seven original top eight P. vivax SEMs, two had minimal levels of genetic diversity uncovered in the bioinformatic analyses (MSP1-19, RAMA), three had similar classification performance (or better using Sal-1) when testing variants (Pv-fam-a, MSP8, RBP2b) and two were not included in this analysis due to being the blacklisted region (EBP, PVX_112670). The latter antigen has since been dropped from the top 8 P. vivax SEMs (Smith et al in preparation) due to other stronger alternatives as the overall assay has been optimized^32^. A recent study has provided evidence of limited natural genetic diversity for P. vivax EBP^42^. From the remaining P. vivax SEMs, there is redundancy in choice of antigens included in the final algorithm^12^. This enables the genetic diversity analysis presented here to guide selection or inclusion of replacement antigens. Both RBP2a and DBPII had high levels of genetic diversity and exhibited differences in the superior construct across two discrete populations (Brazil, Thailand) and are thus not recommended to be included in the final algorithm, although these differences can be overcome when moving from assessment of a single antigen to multiple antigens (i.e. RBP2a Figure 6).

The bioinformatically observed hypervariability and strong selection pressure within DBPII, especially in region II, is consistent with previous investigations^18,43,44^. Earlier reports showed that the region II of DBP serves as a critical domain necessary for P. vivax Duffy binding ligand and DARC interaction^44–46^, and the clusters of polymorphisms opposite the DARC recognition site, called the DEK epitope, are possibly maintained by immune selection pressure^37^. This may be an important finding in the context of developing a P. vivax vaccine targeting DBPII, however, DBPII was not the strongest SEM antigen in terms of individual classification^12^ and the current work provides further evidence to not progress DBPII as a P. vivax SEM. Conversely, whilst RBP2a displayed overall low nucleotide diversity, high haplotype diversity was evident, suggesting the circulation of very rare haplotypes in the population due to the generation of novel polymorphisms during recombination events. There were also prominent signals of variation and balancing selection localised within regions targeted by host immunity^47–49^, consistent with higher levels of diversity observed in previous studies^48,50,51^. These patterns identified bioinformatically were also similar for the antigen RBP2b, however the variant haplotypes selected for RBP2b that were tested immunologically had only few amino acid differences compared to Sal-1 (<13) and this likely resulted in the minimal impact we observed on immunogenicity and classification performance. RBP2a also had few mutational differences to Sal-1 (<8), and no evidence of geographical structuring of diversity, yet resulted in differing immunogenicity and classification performance – demonstrating that it remains essential to immunologically test the impact of antigen sequence diversity.

Another possible P. vivax SEM antigen that could be used to replace MSP3a is MSP5. For MSP5, exon 1 exhibits higher levels of nucleotide diversity than the largely conserved exon 2, which is consistent with prior research^52–54^. Additionally, the observed positive signals of Tajima’s D in exon 1 could indicate immune selection pressure that is maintaining diversity in the population. Interestingly, the variant haplotypes of P. vivax MSP5 tested had the highest number of mutation differences to Sal-1 (29 and 31), however, the immunological results presented here demonstrate no impact on IgG magnitude and classification performance. Along with MSP5, other additional P. vivax SEM antigens assessed here that could still be considered for inclusion in a final panel of eight antigens are S16 (conserved, in line with a prior report^55^, not tested immunologically), RiPR and PTEX150. Both RiPR and PTEX150 had high haplotype diversity, with PTEX150 being geographically structured, but low numbers of mutational differences to Sal-1 and no impact of the variants identified on immunogenicity and classification performance. Furthermore, for all antigens, the P. vivax SEMs likely target multiple epitopes across the protein sequence and this polyclonal response likely also helps overcome the diversity identified.

Of the relatively conserved antigens in the original panel of top eight (MSP1_19_, RAMA^56,57^, Pv-fam-a, MSP8^58^), where we identified limited impact on immunogenicity for those with haplotype variants tested, most had not previously been well-characterized in terms of antigen genetic diversity except MSP1_19_. This MSP1 C-terminal region fragment is genetically conserved, except for the reported E1692K (amino acid position 1709 in previous versions) substitution in previous investigations^59,60^. Interestingly, this amino acid substitution was also observed in the present study but only limited to Asian isolates. Despite having no positive peaks of Tajima’s D detected across the MSP1_19_ sequence (Figure S5), various studies have reported its high immunogenicity in malaria-exposed individuals, immunized animal models, and in-vitro experiments^61–63^, and it has consistently been the 2^nd^ best performing P. vivax SEM (after RBP2b) across studies^12,15^.

In summary, this study comprehensively analysed the genetic diversity and patterns of selection pressure among P. vivax SEMs, and designed and tested haplotype variants more representative of the global sequence population. These experiments have informed inclusion and exclusion of antigens to be used in the multi-antigen P. vivax serological tool. Limitations of this work are that we did not systematically test every single haplotype identified, but rather developed a pipeline to select 1-2 per antigen. We have also not yet experimentally validated the use of a multivalent antigen construct, for antigens such as RBP2a where the top performing construct differed between the two countries tested. Overall, highly diverse antigens can still make attractive targets for developing multivalent vaccine and sero-surveillance markers, and our study demonstrates the importance of testing any identified diversity immunologically. For vaccines, these antigens can be good targets as the immune pressure signifies a potentially important functional role, and multivalent vaccines can be designed to cover the genetic diversity observed in the population, resulting in a broader inhibitory response^18,64^.

## Acknowledgments

We thank all participants and study teams who were involved in the original observational cohorts in Thailand and Brazil. We thank Eamon Conway (WEHI) for assistance with trouble shooting R-code.

## Data sharing statement

Data and code generated for this study are available at: https://github.com/paolobareng/Pvivax_SEM and https://github.com/Longley-Lab/Pv_genetic_diversity_ab_response.

## REFERENCES

1. World Health Organisation. World malaria report 2024: Addressing inequity in the global malaria response. (2024).

2. Robinson, L. J. et al. Strategies for Understanding and Reducing the Plasmodium vivax and Plasmodium ovale Hypnozoite Reservoir in Papua New Guinean Children: A Randomised Placebo-Controlled Trial and Mathematical Model. PLoS Med 12, e1001891 (2015).

3. Commons, R. J., Simpson, J. A., Watson, J., White, N. J. & Price, R. N. Estimating the Proportion of Plasmodium vivax Recurrences Caused by Relapse: A Systematic Review and Meta-Analysis. The American Journal of Tropical Medicine and Hygiene 103, 1094– 1099 (2020).

4. Price, R. N., Commons, R. J., Battle, K. E., Thriemer, K. & Mendis, K. Plasmodium vivax in the Era of the Shrinking P. falciparum Map. Trends in Parasitology 36, 560–570 (2020).

5. Sattabongkot, J., Tsuboi, T., Zollner, G. E., Sirichaisinthop, J. & Cui, L. Plasmodium vivax transmission: chances for control? Trends in Parasitology 20, 192–198 (2004).

6. Hsiang, M. S. et al. Mass drug administration for the control and elimination of Plasmodium vivax malaria: an ecological study from Jiangsu province, China. Malar J 12, 383 (2013).

7. Newby, G. et al. Review of Mass Drug Administration for Malaria and Its Operational Challenges. The American Journal of Tropical Medicine and Hygiene 93, 125–134 (2015).

8. Alonso, P. L. The Role of Mass Drug Administration of Antimalarials. The American Journal of Tropical Medicine and Hygiene 103, 1–2 (2020).

9. Ruwende, C. & Hill, A. Glucose-6-phosphate dehydrogenase deficiency and malaria. J Mol Med 76, 581–588 (1998).

10. Varo, R., Balanza, N., Mayor, A. & Bassat, Q. Diagnosis of clinical malaria in endemic settings. Expert Review of Anti-infective Therapy 19, 79–92 (2021).

11. Kim, S., Luande, V. N., Rocklöv, J., Carlton, J. M. & Tozan, Y. A Systematic Review of the Evidence on the Effectiveness and Cost-Effectiveness of Mass Screen-and-Treat Interventions for Malaria Control. The American Journal of Tropical Medicine and Hygiene 105, 1722–1731 (2021).

12. Longley, R. J. et al. Development and validation of serological markers for detecting recent Plasmodium vivax infection. Nature Medicine 26, 741–749 (2020).

13. Kartal, L., Mueller, I. & Longley, R. J. Using Serological Markers for the Surveillance of Plasmodium vivax Malaria: A Scoping Review. Pathogens 12, 791 (2023).

14. White, N. J. Determinants of relapse periodicity in Plasmodium vivax malaria. Malar J 10, 297 (2011).

15. Tayipto, Y., Liu, Z., Mueller, I. & Longley, R. J. Serology for Plasmodium vivax surveillance: A novel approach to accelerate towards elimination. Parasitology International 87, 102492 (2022).

16. Greenhouse, B. et al. Priority use cases for antibody-detecting assays of recent malaria exposure as tools to achieve and sustain malaria elimination. Gates Open Res 3, 131 (2019).

17. Nekkab, N. et al. Accelerating towards P. vivax elimination with a novel serological test-and-treat strategy: a modelling case study in Brazil. Lancet Reg Health Am 22, 100511 (2023).

18. de Sousa, T. N., Carvalho, L. H. & de Brito, C. F. A. Worldwide genetic variability of the duffy binding protein: Insights into Plasmodium vivax vaccine development. PLoS ONE 6, e22944–e22944 (2011).

19. Arnott, A. et al. Global Population Structure of the Genes Encoding the Malaria Vaccine Candidate, Plasmodium vivax Apical Membrane Antigen 1 (PvAMA1). PLoS Neglected Tropical Diseases 7, (2013).

20. Barry, A. E. & Arnott, A. Strategies for designing and monitoring malaria vaccines targeting diverse antigens. Frontiers in Immunology 5, (2014).

21. Naung, M. T. et al. Global diversity and balancing selection of 23 leading Plasmodium falciparum candidate vaccine antigens. PLoS Comput Biol 18, e1009801 (2022).

22. Pearson, R. D. et al. Genomic analysis of local variation and recent evolution in Plasmodium vivax. Nature Genetics 48, 959–964 (2016).

23. Hupalo, D. N. et al. Population genomics studies identify signatures of global dispersal and drug resistance in Plasmodium vivax. Nature Genetics 48, 953–958 (2016).

24. Longley, R. J. et al. Plasmodium vivax malaria serological exposure markers: Assessing the degree and implications of cross-reactivity with P. knowlesi. Cell Reports Medicine 3, 100662 (2022).

25. Auburn, S. et al. A new Plasmodium vivax reference sequence with improved assembly of the subtelomeres reveals an abundance of pir genes. Wellcome Open Res 1, 4 (2016).

26. Van der Auwera, G. A. et al. From fastQ data to high-confidence variant calls: The genome analysis toolkit best practices pipeline. Current Protocols in Bioinformatics (2013).

27. Lee S, et al. Assessing clonality in malaria parasites using massively parallel sequencing data. in preparation (2016).

28. Leigh, J. W. & Bryant, D. POPART: full-feature software for haplotype network construction. researchgate.net (2015) doi:10.1111/2041-210X.12410.

29. Jumper, J. et al. Highly accurate protein structure prediction with AlphaFold. Nature 596, 583–589 (2021).

30. Schrödinger, LLC. The PyMOL Molecular Graphics System, Version 2.0.

31. Longley, R. J. et al. Naturally acquired antibody responses to more than 300 Plasmodium vivax proteins in three geographic regions. PLoS Negl Trop Dis 11, e0005888 (2017).

32. Mazhari, R. et al. A comparison of non-magnetic and magnetic beads for measuring IgG antibodies against Plasmodium vivax antigens in a multiplexed bead-based assay using Luminex technology (Bio-Plex 200 or MAGPIX). PLoS ONE 15, e0238010 (2020).

33. Nguitragool, W. et al. Highly heterogeneous residual malaria risk in western Thailand. International Journal for Parasitology 49, 455–462 (2019).

34. Monteiro, W. et al. Prevalence and force of Plasmodium vivax blood-stage infection and associated clinical malaria burden in the Brazilian Amazon. Mem Inst Oswaldo Cruz 117, e210330 (2022).

35. Robin, X. et al. pROC: an open-source package for R and S+ to analyze and compare ROC curves. BMC Bioinformatics 12, 77 (2011).

36. Wright, M. N. & Ziegler, A. ranger: A Fast Implementation of Random Forests for High Dimensional Data in C++ and R. J. Stat. Soft. 77, (2017).

37. Batchelor, J. D. et al. Red Blood Cell Invasion by Plasmodium vivax: Structural Basis for DBP Engagement of DARC. PLoS Pathog 10, e1003869 (2014).

38. Obadia, T. et al. Developing sero-diagnostic tests to facilitate Plasmodium vivax Serological Test-and-Treat approaches: modeling the balance between public health impact and overtreatment. BMC Med 20, 98 (2022).

39. World Health Organisation. Malaria vaccines: preferred product characteristics and clinical development considerations.

40. Rosado, J. et al. Malaria transmission structure in the Peruvian Amazon through antibody signatures to Plasmodium vivax. PLoS Negl Trop Dis 16, e0010415 (2022).

41. Grimée, M. et al. Using serological diagnostics to characterize remaining high-incidence pockets of malaria in forest-fringe Cambodia. Malar J 23, 49 (2024).

42. Fernandes, G. M. et al. Natural genetic diversity of the DBL domain of a novel member of the Plasmodium vivax erythrocyte binding-like proteins (EBP2) in the Amazon rainforest. Infection, Genetics and Evolution 123, 105628 (2024).

43. Singh, S. K., Hora, R., Belrhali, H., Chitnis, C. E. & Sharma, A. Structural basis for Duffy recognition by the malaria parasite Duffy-binding-like domain. Nature (2006).

44. Xainli, J., Adams, J. H. & King, C. L. The erythrocyte binding motif of Plasmodium vivax Duffy binding protein is highly polymorphic and functionally conserved in isolates from Papua New Guinea. Molecular and Biochemical Parasitology 111, 253–260 (2000).

45. Tsuboi,’, Takafumi et al. Natural Variation within the Principal Adhesion Domain of the Plasmodium vivax Duffy Binding Protein. Infection and Immunity 180, 497–506 (1994).

46. Baum, J., Thomas, A. W. & Conway, D. J. Evidence for Diversifying Selection on Erythrocyte-Binding Antigens of Plasmodium falciparum and P. vivax.

47. Gruszczyk, J. et al. Transferrin receptor 1 is a reticulocyte-specific receptor for *Plasmodium vivax*. Science 359, 48–55 (2018).

48. Gruszczyk, J. et al. Structurally conserved erythrocyte-binding domain in *Plasmodium* provides a versatile scaffold for alternate receptor engagement. Proc. Natl. Acad. Sci. U.S.A. 113, (2016).

49. Gruszczyk, J. et al. Cryo-EM structure of an essential Plasmodium vivax invasion complex. Nature 559, 135–139 (2018).

50. Zhang, X. et al. Genetic diversity of Plasmodium vivax reticulocyte binding protein 2b in global parasite populations. Parasites Vectors 15, 205 (2022).

51. Kosaisavee, V. et al. Genetic diversity in new members of the reticulocyte binding protein family in Thai Plasmodium vivax isolates. PLoS ONE 7, (2012).

52. Putaporntip, C., Udomsangpetch, R., Pattanawong, U., Cui, L. & Jongwutiwes, S. Genetic diversity of the Plasmodium vivax merozoite surface protein-5 locus from diverse geographic origins. Gene 456, 24–35 (2010).

53. Black, C. G., Barnwell, J. W., Huber, C. S., Galinski, M. R. & Coppel, R. L. The Plasmodium vivax homologues of merozoite surface proteins 4 and 5 from Plasmodium falciparum are expressed at different locations in the merozoite. Molecular and Biochemical Parasitology 120, 215–224 (2002).

54. Gomez, A., Suarez, C. F., Martinez, P., Saravia, C. & Patarroyo, M. A. High polymorphism in Plasmodium vivax merozoite surface protein-5 (MSP5). Parasitology 133, 661 (2006).

55. Ford, A. et al. Gene Polymorphisms Among Plasmodium vivax Geographical Isolates and the Potential as New Biomarkers for Gametocyte Detection. Front. Cell. Infect. Microbiol. 11, 789417 (2022).

56. Lu, F. et al. Profiling the humoral immune responses to Plasmodium vivax infection and identification of candidate immunogenic rhoptry-associated membrane antigen (RAMA). Journal of Proteomics 102, 66–82 (2014).

57. Ge, J. et al. Immunogenicity and antigenicity of a conserved fragment of the rhoptry-associated membrane antigen of Plasmodium vivax. Parasites Vectors 15, 428 (2022).

58. Pacheco, M. A. et al. Evidence of purifying selection on merozoite surface protein 8 (MSP8) and 10 (MSP10) in Plasmodium spp. Infection, Genetics and Evolution 12, 978– 986 (2012).

59. Pasay, Ma. C., Cheng, Q., Rzepczyk, C. & Saul, A. Dimorphism of the C terminus of the Plasmodium vivax merozoite surface protein 1. Molecular and Biochemical Parasitology 70, 217–219 (1995).

60. Putaporntip, C. et al. Mosaic organization and heterogeneity in frequency of allelic recombination of the Plasmodium vivax merozoite surface protein-1 locus. Proceedings of the National Academy of Sciences 99, 16348–16353 (2002).

61. Soares, I. S. et al. A Plasmodium vivax vaccine candidate displays limited allele polymorphism, which does not restrict recognition by antibodies. Molecular Medicine 5, 459–470 (1999).

62. María Espinosa, A., et al. Expression, polymorphism analysis, reticulocyte binding and serological reactivity of two Plasmodium vivax MSP-1 protein recombinant fragments. Vaccine 21, 1033–1043 (2003).

63. Min, H. M. K. et al. Immunogenicity of the Plasmodium vivax merozoite surface protein 1 paralog in the induction of naturally acquired antibody and memory B cell responses. Malaria Journal 16, 354–354 (2017).

64. Duan, J. et al. Population structure of the genes encoding the polymorphic *Plasmodium falciparum* apical membrane antigen 1: Implications for vaccine design. Proc. Natl. Acad. Sci. U.S.A. 105, 7857–7862 (2008).

