## Supplemental Tables and Figures for "Navigating parasite antigen genetic diversity in the design of Plasmodium vivax serological exposure markers for malaria"

### BARENG, WU ET AL SUPPLEMENTARY MATERIAL

**Contents**

Figure S1. Pipeline for antigen inclusion/exclusion.

Figure S2. SDS-gels of *P. vivax* WGCF-expressed variant proteins.

Figure S3. Line plots showing nucleotide diversity calculated in a sliding window approach for all 11 *P. vivax* antigens across different populations

Figure S4. Spatial diversity displayed over the modelled three-dimensional protein structures for DBPII, RBP2a and RBP2b.

Figure S5. Line plots showing Tajima's D calculated in a sliding window approach for all 11 *P. vivax* antigens across different populations.

Figure S6. Important gene domains or sites mapped over three-dimensional protein structures.

Figure S7. Correlation of IgG antibody levels to all *P. vivax* serological exposure marker antigen constructs in (A) Brazil and (B) Thailand.

Figure S8. ROC curves comparing classification performance of *P. vivax* serological exposure markers when using the reference Sal-1 strain compared to identified variant haplotypes.

Table S1. Summary results of diversity measures and Tajima's D calculated for 11 *P. vivax* antigens across different populations.

Table S2. *P. vivax* reference (Sal-1) and variant (haplotype) proteins tested in immunogenicity experiments.

Table S3. AUC values for each *P. vivax* serological exposure marker construct tested.

Table S4. Top antigen combinations (ranging from two to eight) for the combined dataset (Brazil, Thailand and Negative Controls), as well as the Brazil and Thailand cohort datasets.

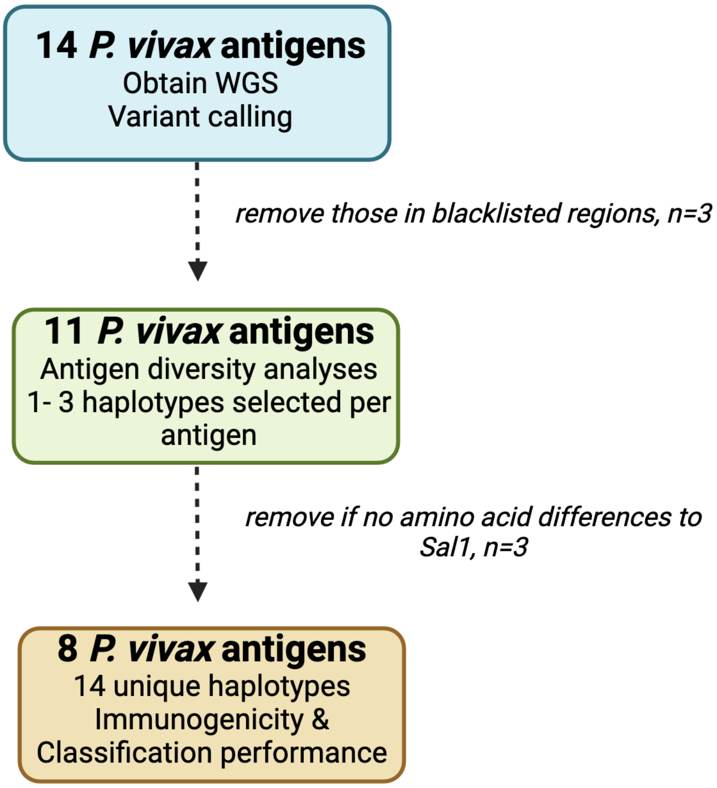


**Figure S1.** Pipeline for antigen inclusion/exclusion.

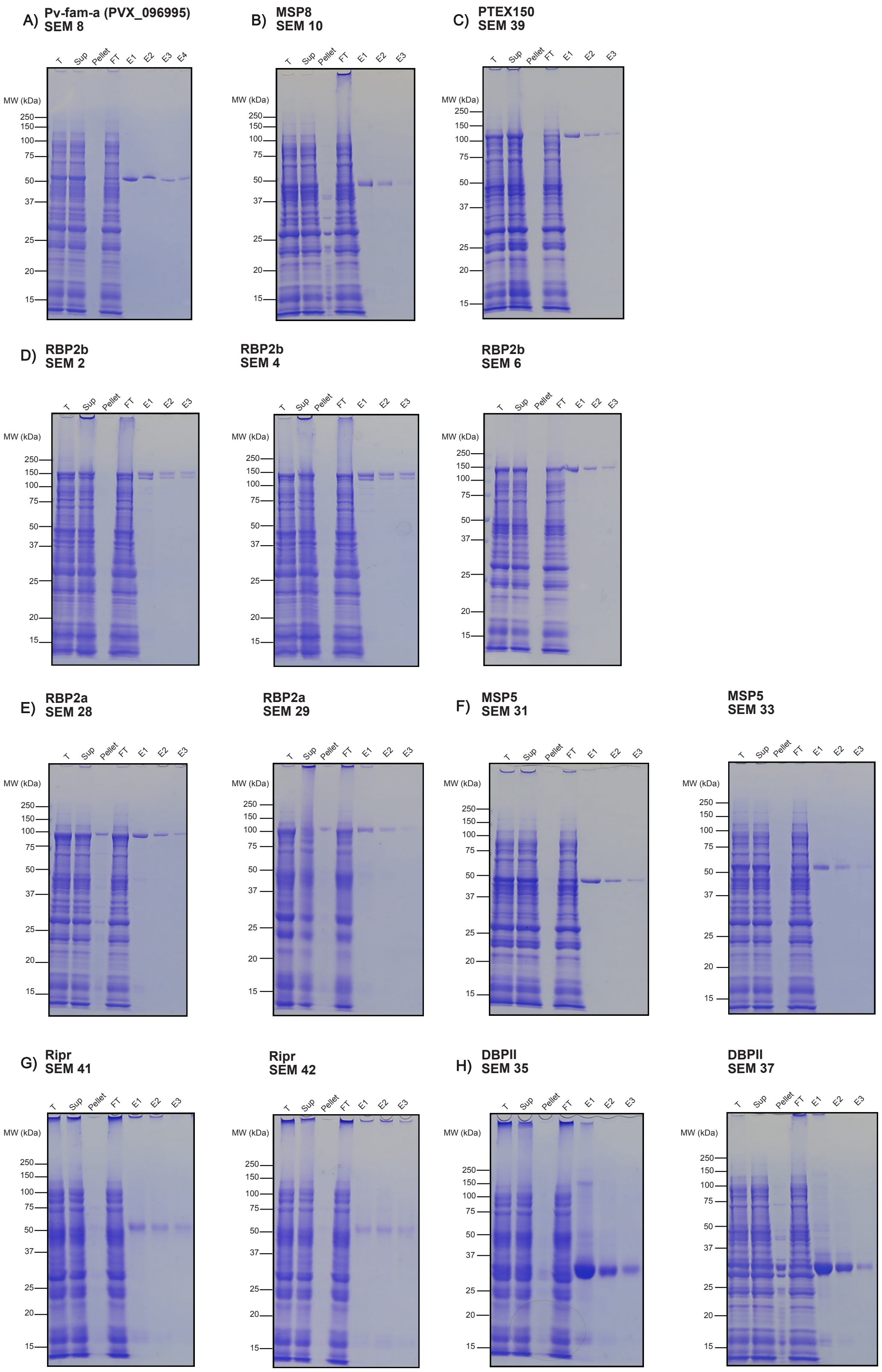


**Figure S2.** SDS-PAGE with 12.5% gels of *P. vivax* WGCF-expressed variant proteins, stained with Coomassie Brilliant Blue (CBB). The total reaction mixture was centrifuged prior to loading onto the affinity column. **T** refers to the total reaction mixture; **Sup** and **Pellet** indicate the supernatant and pellet of **T**; **FT** is the flow-through from the affinity purification column; **E1–E3** are elution fractions. Predicted MW (kDa) with His-tag were as follows: A) Pv-fam-a$_{61\text{-}421}$ 48.92, B) MSP8$_{24\text{-}463}$ 50.27, C) PTEX150$_{24\text{-}408}$ 97.36, D) RBP2b$_{161\text{-}1454}$ all 152.7, E) RBP2a$_{160\text{-}1135}$ all 114.52, F) MSP5$_{23\text{-}365}$ SEM31 37.23, SEM33 37.54, G) RIPR$_{552\text{-}1075}$ both 59.46, H) DBPII$_{242\text{-}529}$ SEM35 34.84, SEM37 34.71.

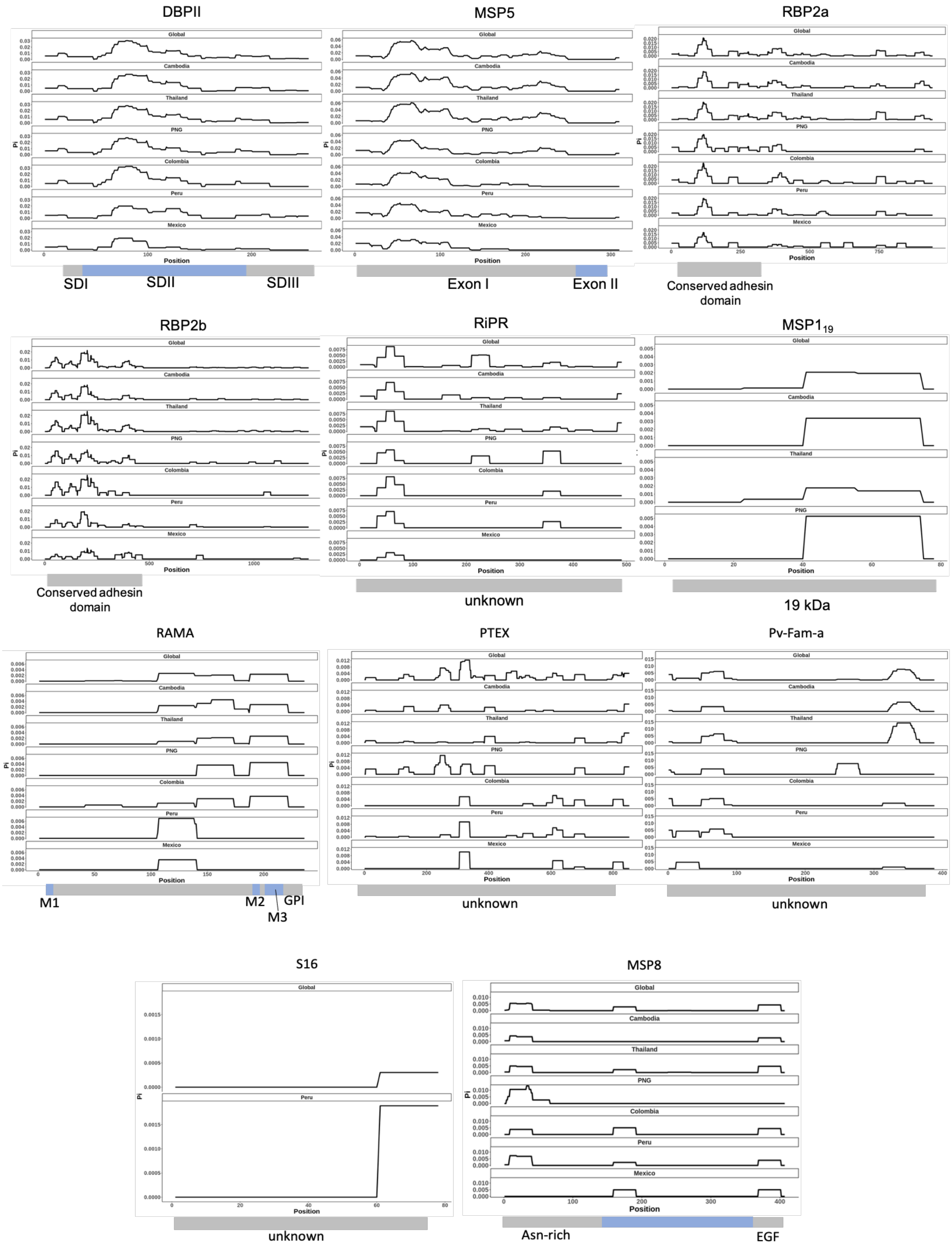


**Figure S3.** Line plots showing nucleotide diversity calculated in a sliding window approach for all 11 *P. vivax* antigens across different populations. Note the y-axis varies between antigens. The coloured blocks below each plot display the known protein domains based from previous studies, with some being unknown.

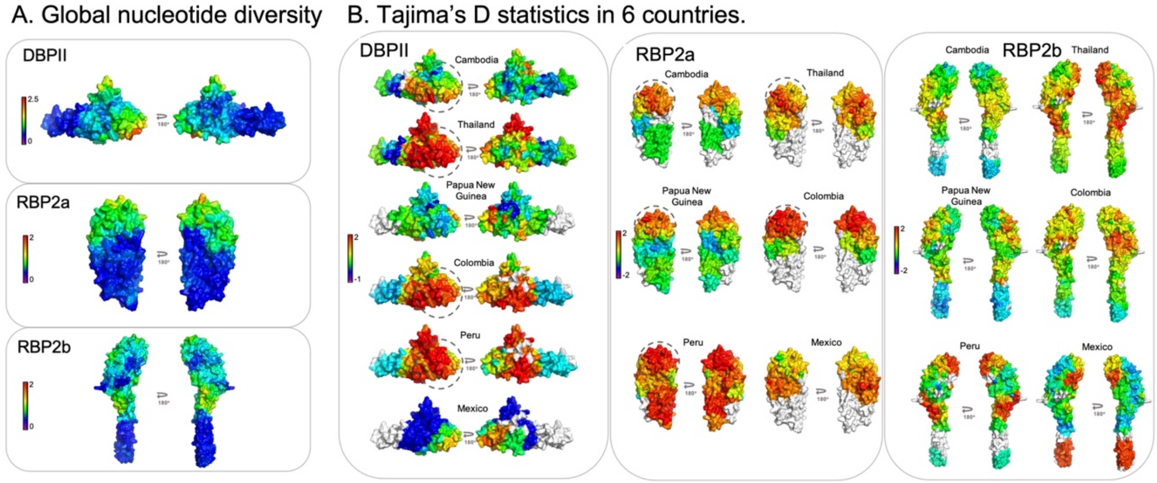


**Figure S4.** Spatial diversity displayed over the modelled three-dimensional protein structures for DBPII, RBP2a and RBP2b. Crystal structures 4Z8N (RBP2a); 5W53 (RPB2b); 4Y5S (DBPII) were used as references**.** (A) Global nucleotide diversity. The colour spectrum bar shows the π values, with red being the highest possible value for diversity while blue indicates no diversity. (B) Tajima's D statistics. The legend in the colour spectrum shows the Tajima's D values, with red indicating high positive selection and blue indicating negative selection pressure.

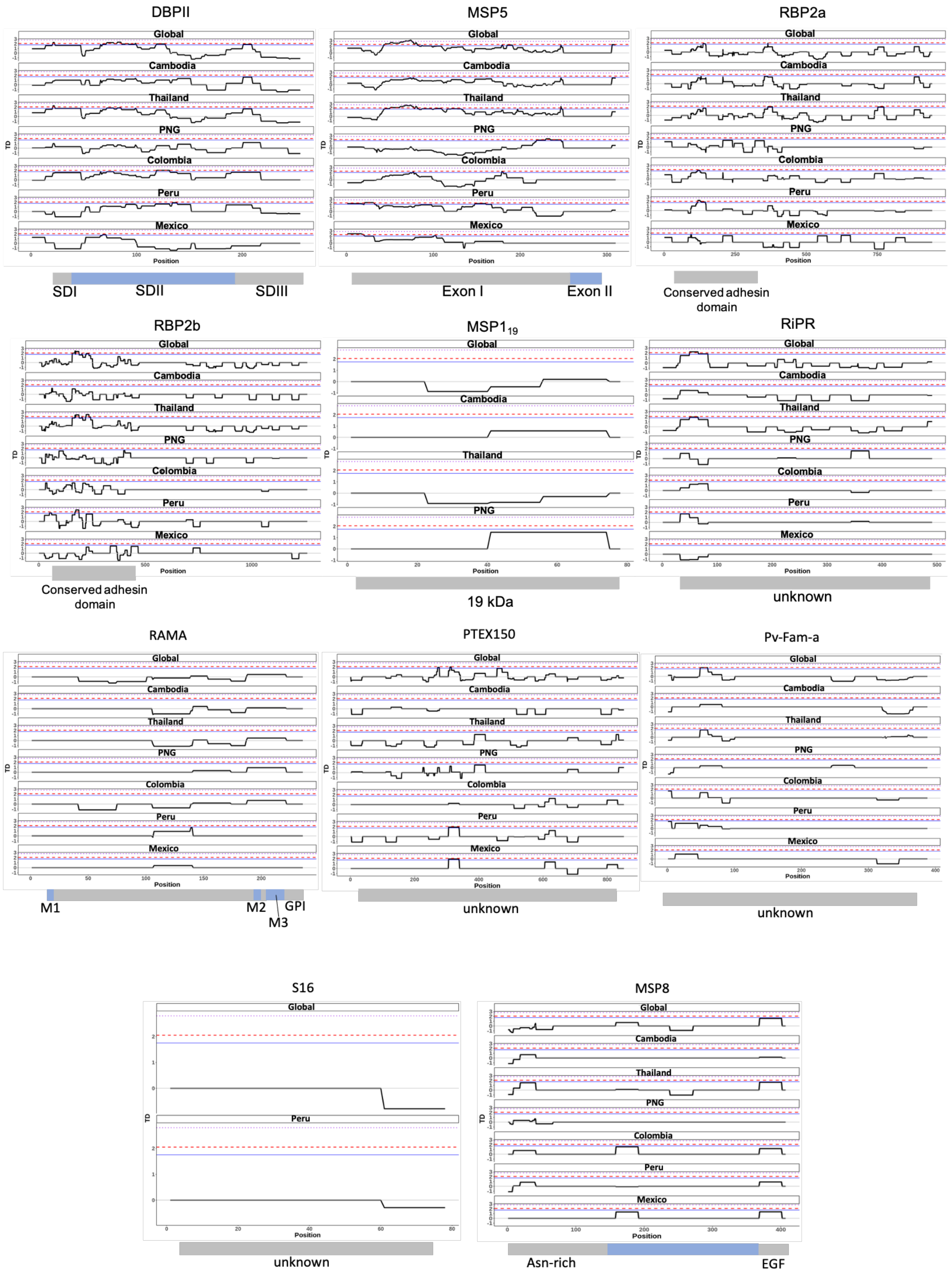


**Figure S5.** Line plots showing Tajima's D calculated in a sliding window approach for all 11 *P. vivax* antigens across different populations. The blue square dot lines indicate p<0.01; red dashed lines indicate p<0.05; blue solid lines represent p<0.10.

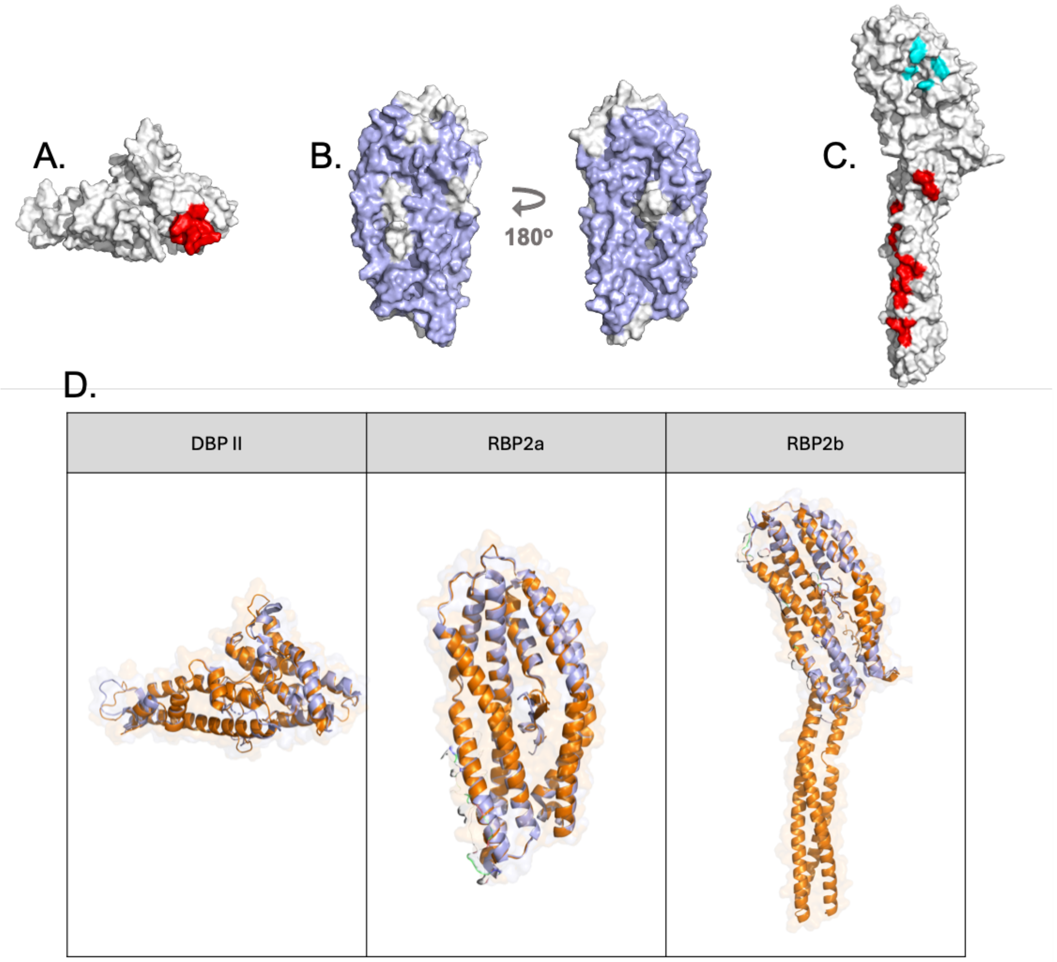


**Figure S6.** Important gene domains or sites mapped over three-dimensional protein structures. (A) DBPII. The highly polymorphic DEK epitope (Sal-1: DEKAQQRRKQ; PvP01: GEKAQQHRKQ) highlighted in red. (B) RBP2a. The conserved adhesin domain highlighted in light blue. (C) RBP2b. C-terminal: TfR1 binding site (red) and N-terminal: Tf binding site (cyan). (D) Structural comparison of DBPII, RBP2a, and RBP2b protein models showing high similarity in the overlapping regions. Light blue regions represent experimentally determined structures, while orange regions correspond to AlphaFold2 predictions. Across all three proteins, the root mean square deviation (RMSD) for the aligned regions is below 1 Å, indicating strong structural agreement between experimental and predicted models.

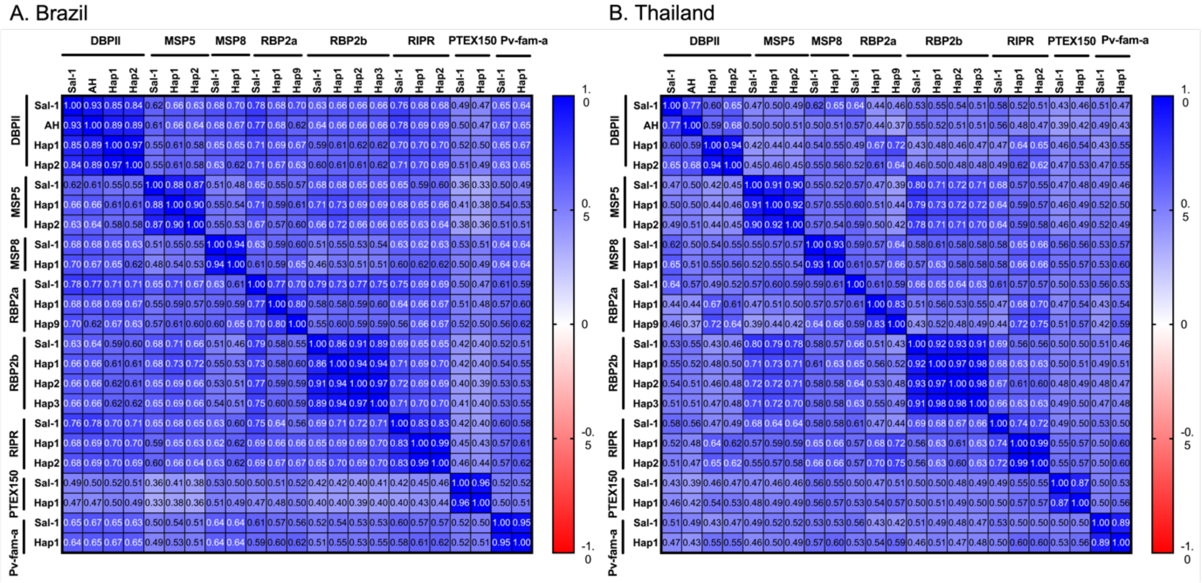


**Figure S7.** Correlation of IgG antibody levels to all *P. vivax* serological exposure marker antigen constructs in (A) Brazil and (B) Thailand. Spearman r values are shown. Antigens are in the same order as listed on the horizontal titles.

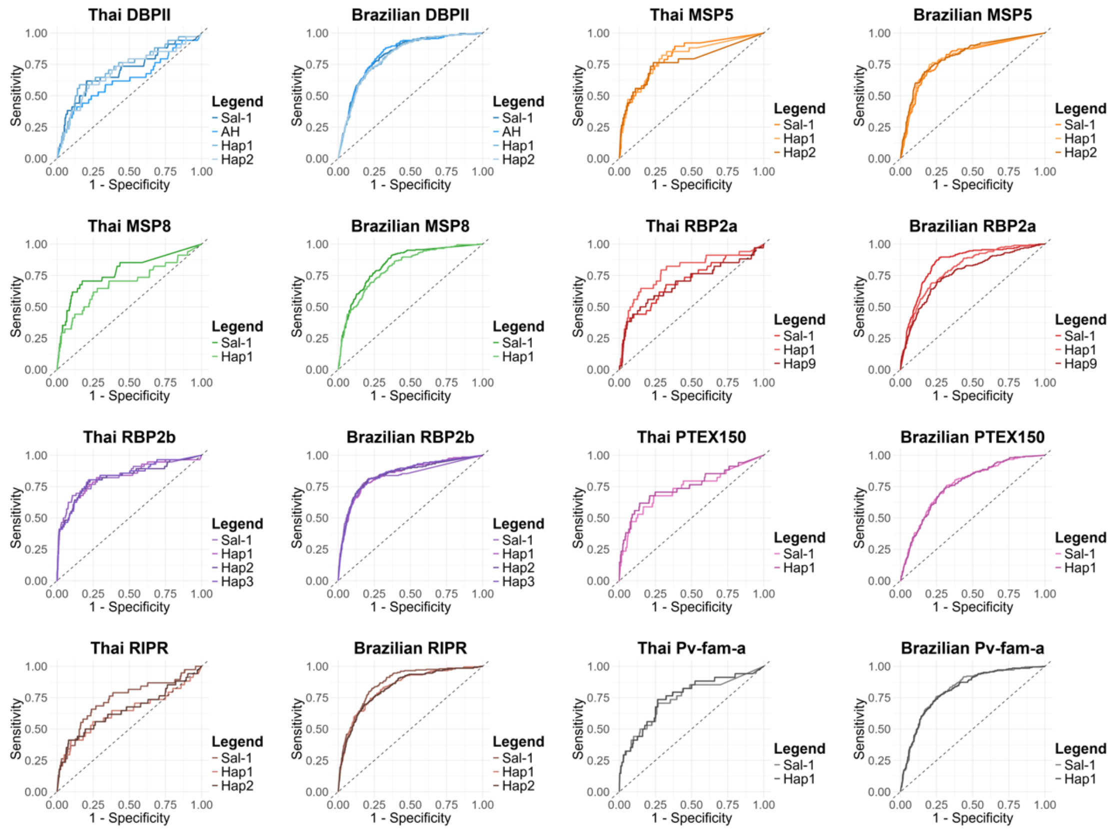


**Figure S8.** ROC curves comparing classification performance of *P. vivax* serological exposure markers when using the reference Sal-1 strain compared to identified variant haplotypes.

**Table S1.** Summary results of diversity measures and Tajima's D calculated for 11 *P. vivax* antigens across different populations. # seq: number of sequences; SNP: Single Nucleotide Polymorphisms; πx10^-3: Nucleotide Diversity; NSP: Non-Synonymous polymorphisms; SP: Synonymous polymorphisms; # Hap: number of haplotypes; Hd: Haplotype diversity. Amino acid region covered is shown within the antigen title, with sequenes mapped using P01 as the reference.

| Antigen | Country | # seq | Nucleotide sequence length | SNPs | $\pi x10^{-3}$ | Tajima 's D | NSP | SP | # Hap | Hd |
|---|---|---|---|---|---|---|---|---|---|---|
| *dbpII* (aa 242-529) | **Global** | **278** | **864** | **30** | **7.652** | **1.008** | **27** | **3** | **65** | **0.957** |
| | Thailand | 109 | 864 | 26 | 7.289 | 0.815 | 26 | 0 | 43 | 0.96 |
| | Myanmar | 9 | 864 | 14 | 6.655 | 0.558 | 14 | 0 | 7 | 0.944 |
| | Cambodia | 29 | 864 | 20 | 6.979 | 0.645 | 20 | 0 | 16 | 0.946 |
| | Vietnam | 9 | 864 | 17 | 8.166 | 0.623 | 17 | 0 | 8 | 0.972 |
| | Malaysia | 6 | 864 | 13 | 6.25 | -0.315 | 13 | 0 | 4 | 0.8 |
| | Indonesia | 5 | 864 | 13 | 7.292 | 0.07 | 13 | 0 | 5 | 1 |
| | Colombia | 31 | 864 | 18 | 7.701 | 1.632 | 17 | 1 | 11 | 0.901 |
| | Mexico | 20 | 864 | 12 | 3.801 | -0.104 | 12 | 0 | 5 | 0.558 |
| | Peru | 42 | 864 | 17 | 5.866 | 0.91 | 17 | 0 | 16 | 0.918 |
| | Brazil | 4 | 864 | 14 | 11.767 | 3.361 | 13 | 1 | 3 | 0.833 |
| | PNG | 14 | 864 | 19 | 7.008 | 0.056 | 19 | 0 | 10 | 0.956 |
| *rbp2b* (aa 161-1454) | **Global** | **269** | **3882** | **54** | **2.014** | **-0.158** | **47** | **7** | **133** | **0.985** |
| | Thailand | 104 | 3882 | 46 | 2.187 | 0.094 | 40 | 6 | 72 | 0.985 |
| | Myanmar | 9 | 3882 | 14 | 1.259 | -0.244 | 11 | 3 | 6 | 0.833 |
| | Cambodia | 29 | 3882 | 29 | 1.518 | -0.627 | 26 | 3 | 27 | 0.995 |
| | Vietnam | 10 | 3882 | 24 | 2.295 | 0.455 | 23 | 1 | 9 | 0.978 |
| | Malaysia | 6 | 3882 | 7 | 0.876 | 0.635 | 7 | 0 | 4 | 0.867 |
| | Indonesia | 6 | 3882 | 20 | 2.284 | 0.077 | 19 | 1 | 6 | 1 |
| | Colombia | 30 | 3882 | 34 | 2.124 | -0.037 | 33 | 1 | 23 | 0.97 |
| | Mexico | 19 | 3882 | 21 | 1.399 | -0.37 | 19 | 2 | 5 | 0.696 |
| | Peru | 40 | 3882 | 22 | 1.354 | 0.055 | 18 | 4 | 15 | 0.915 |
| | Brazil | 4 | 3882 | 20 | 3.177 | 1.338 | 19 | 1 | 4 | 1 |
| | PNG | 12 | 3882 | 26 | 1.983 | -0.313 | 23 | 3 | 10 | 0.955 |
| *rbp2a* (aa 160-1135) | **Global** | **281** | **2928** | **49** | **2.297** | **-0.424** | **43** | **6** | **133** | **0.981** |
| | Thailand | 106 | 2928 | 36 | 2.331 | -0.023 | 31 | 5 | 50 | 0.959 |
| | Myanmar | 9 | 2928 | 16 | 2.068 | 0.139 | 16 | 0 | 9 | 1 |
| | Cambodia | 33 | 2928 | 26 | 2.122 | -0.106 | 24 | 2 | 21 | 0.966 |
| | Vietnam | 10 | 2928 | 19 | 2.558 | 0.539 | 18 | 1 | 9 | 0.978 |
| | Malaysia | 6 | 2928 | 10 | 1.753 | 1.034 | 10 | 0 | 6 | 1 |
| | Indonesia | 7 | 2928 | 17 | 2.521 | 0.355 | 16 | 1 | 7 | 1 |
| | Colombia | 31 | 2928 | 24 | 2.435 | 0.658 | 23 | 1 | 18 | 0.959 |
| | Mexico | 20 | 2928 | 16 | 1.65 | 0.265 | 16 | 0 | 8 | 0.774 |
| | Peru | 41 | 2928 | 19 | 1.57 | 0.115 | 19 | 0 | 21 | 0.94 |

| | | | | | | | | | | |
|---|---|---|---|---|---|---|---|---|---|---|
| | Brazil | 4 | 2928 | 5 | 0.968 | 0.372 | 5 | 0 | 3 | 0.833 |
| | PNG | 14 | 2928 | 16 | 1.899 | 0.432 | 16 | 0 | 10 | 0.923 |
| *ripr* (aa 552-1075) | **Global** | **271** | **1572** | **16** | **1.317** | **-0.5** | **8** | **8** | **24** | **0.893** |
| | Thailand | 106 | 1572 | 13 | 1.2 | -0.643 | 7 | 6 | 12 | 0.842 |
| | Myanmar | 9 | 1572 | 8 | 2.05 | 0.43 | 5 | 3 | 6 | 0.917 |
| | Cambodia | 31 | 1572 | 8 | 0.925 | -0.826 | 6 | 2 | 9 | 0.748 |
| | Vietnam | 9 | 1572 | 3 | 0.442 | -1.417 | 2 | 1 | 3 | 0.417 |
| | Malaysia | 6 | 1572 | 4 | 1.145 | 0.149 | 3 | 1 | 5 | 0.933 |
| | Indonesia | 6 | 1572 | 3 | 0.848 | 0.076 | 3 | 0 | 4 | 0.8 |
| | Colombia | 29 | 1572 | 3 | 0.638 | 0.742 | 3 | 0 | 5 | 0.727 |
| | Mexico | 19 | 1572 | 2 | 0.201 | -1.071 | 2 | 0 | 3 | 0.205 |
| | Peru | 39 | 1572 | 3 | 0.616 | 0.802 | 3 | 0 | 5 | 0.698 |
| | Brazil | | | 0 | 0 | 0 | 0 | 0 | 1 | 0 |
| | PNG | 16 | 1572 | 4 | 0.917 | 0.613 | 4 | 0 | 6 | 0.85 |
| *msp5* (aa 23-365) | **Global** | **278** | **1029** | **58** | **14.838** | **2.017** | **53** | **5** | **150** | **0.991** |
| | Thailand | 104 | 1029 | 61 | 16.477 | 1.677 | 56 | 5 | 64 | 0.984 |
| | Myanmar | 9 | 1029 | 43 | 15.522 | 0.048 | 40 | 3 | 9 | 1 |
| | Cambodia | 30 | 1029 | 56 | 15.871 | 0.825 | 52 | 4 | 26 | 0.989 |
| | Vietnam | 10 | 1029 | 48 | 17.406 | 0.495 | 44 | 4 | 10 | 1 |
| | Malaysia | 6 | 1029 | 34 | 15.873 | 1.05 | 33 | 1 | 6 | 1 |
| | Indonesia | 7 | 1029 | 34 | 13.235 | 0.436 | 32 | 2 | 7 | 1 |
| | Colombia | 31 | 1029 | 33 | 8.755 | 0.328 | 31 | 2 | 19 | 0.933 |
| | Mexico | 19 | 1029 | 21 | 8.059 | 1.469 | 20 | 1 | 7 | 0.749 |
| | Brazil | 4 | 1029 | 12 | 6.641 | 0.444 | 12 | 0 | 4 | 1 |
| | Peru | 42 | 1029 | 35 | 11.193 | 1.438 | 33 | 2 | 25 | 0.963 |
| | PNG | 16 | 1029 | 40 | 10.965 | -0.168 | 38 | 2 | 14 | 0.983 |
| *msp8* (aa 24-463) | **Global** | **266** | **1320** | **9** | **0.987** | **-0.239** | **8** | **1** | **10** | **0.641** |
| | Thailand | 98 | 1320 | 5 | 0.909 | 0.506 | 4 | 1 | 7 | 0.59 |
| | Myanmar | 9 | 1320 | 4 | 1.326 | 0.768 | 3 | 1 | 4 | 0.583 |
| | Cambodia | 31 | 1320 | 3 | 0.536 | -0.135 | 2 | 1 | 3 | 0.411 |
| | Vietnam | 9 | 1320 | 2 | 0.673 | 0.715 | 2 | 0 | 3 | 0.722 |
| | Malaysia | 6 | 1320 | 1 | 0.303 | -0.338 | 0 | 1 | 1 | 0 |
| | Indonesia | 6 | 1320 | 2 | 0.758 | 0.671 | 1 | 1 | 2 | 0.6 |
| | Colombia | 29 | 1320 | 3 | 1.006 | 1.753 | 2 | 1 | 3 | 0.675 |
| | Mexico | 19 | 1320 | 2 | 0.749 | 1.738 | 1 | 1 | 2 | 0.491 |
| | Peru | 40 | 1320 | 5 | 1.016 | 0.358 | 4 | 1 | 5 | 0.696 |
| | Brazil | 4 | 1320 | 2 | 0.884 | 0.592 | 1 | 1 | 2 | 0.5 |
| | PNG | 14 | 1320 | 4 | 0.999 | 0.159 | 4 | 0 | 4 | 0.747 |
| *rama* (aa 462-730) | **Global** | **271** | **807** | **7** | **0.927** | **-0.696** | **4** | **3** | **5** | **0.467** |
| | Thailand | 105 | 807 | 5 | 0.732 | -0.808 | 3 | 2 | 4 | 0.351 |
| | Myanmar | 9 | 807 | 1 | 0.31 | -0.881 | 0 | 1 | 1 | 0 |

| | | | | | | | | | | |
|---|---|---|---|---|---|---|---|---|---|---|
| | Cambodia | 31 | 807 | 5 | 1.205 | -0.604 | 3 | 2 | 4 | 0.475 |
| | Vietnam | 10 | 807 | 3 | 0.991 | -0.884 | 1 | 2 | 2 | 0.356 |
| | Malaysia | 6 | 807 | 1 | 0.661 | 0.851 | 1 | 0 | 1 | 0 |
| | Indonesia | 4 | 807 | 1 | 0.826 | 1.633 | 1 | 0 | 1 | 0 |
| | Colombia | 30 | 807 | 4 | 1.06 | -0.393 | 2 | 2 | 3 | 0.467 |
| | Mexico | 19 | 807 | 1 | 0.435 | 0.417 | 1 | 0 | 1 | 0 |
| | Peru | 39 | 807 | 2 | 0.861 | 0.892 | 2 | 0 | 3 | 0.602 |
| | Brazil | 4 | 807 | 2 | 1.239 | -0.71 | 1 | 1 | 2 | 0.5 |
| | PNG | 14 | 807 | 2 | 1.008 | 0.795 | 1 | 1 | 2 | 0.44 |
| *ptex150* (aa 24-908) | **Global** | **267** | **2655** | **37** | **1.972** | **-0.142** | **27** | **10** | **57** | **0.9** |
| | Thailand | 102 | 2655 | 15 | 0.665 | -1.071 | 13 | 2 | 16 | 0.699 |
| | Myanmar | 9 | 2655 | 4 | 0.45 | -0.766 | 3 | 1 | 4 | 0.694 |
| | Cambodia | 28 | 2655 | 10 | 0.64 | -1.091 | 8 | 2 | 7 | 0.796 |
| | Vietnam | 9 | 2655 | 4 | 0.345 | -1.533 | 3 | 1 | 4 | 0.583 |
| | Malaysia | 6 | 2655 | 4 | 0.703 | 0.355 | 4 | 0 | 5 | 0.933 |
| | Indonesia | 6 | 2655 | 8 | 1.532 | 0.947 | 8 | 0 | 5 | 0.933 |
| | Colombia | 31 | 2655 | 8 | 0.781 | 0.106 | 5 | 3 | 7 | 0.744 |
| | Mexico | 19 | 2655 | 5 | 0.767 | 1.314 | 3 | 2 | 3 | 0.69 |
| | Peru | 39 | 2655 | 11 | 0.867 | -0.351 | 8 | 3 | 6 | 0.659 |
| | Brazil | 4 | 2655 | 7 | 1.758 | 2.18 | 6 | 1 | 3 | 0.833 |
| | PNG | 16 | 2655 | 12 | 1.491 | 0.361 | 11 | 1 | 13 | 0.967 |
| $MSP1_{19}$ (aa 1605-1712) | **Global** | **286** | **333** | **2** | **0.624** | **-0.456** | **1** | **1** | **2** | **0.194** |
| | Thailand | 107 | 333 | 2 | 0.531 | -0.815 | 1 | 1 | 2 | 0.14 |
| | Myanmar | 36 | 333 | 2 | 0.777 | -0.901 | 1 | 1 | 2 | 0.203 |
| | Cambodia | 34 | 333 | 1 | 1.017 | 0.576 | 1 | 0 | 1 | 0 |
| | Vietnam | 11 | 333 | 0 | 0 | NA | 0 | 0 | 1 | 0 |
| | Indonesia | 7 | 333 | 1 | 1.43 | 0.559 | 1 | 0 | 1 | 0 |
| | Malaysia | 6 | 333 | 0 | 0 | NA | 0 | 0 | 1 | 0 |
| | Colombia | 31 | 333 | 0 | 0 | NA | 0 | 0 | 1 | 0 |
| | Mexico | 19 | 333 | 0 | 0 | NA | 0 | 0 | 1 | 0 |
| | Brazil | 4 | 333 | 0 | 0 | NA | 0 | 0 | 1 | 0 |
| *s16* (aa 31-140) | **Global** | **261** | **333** | **1** | **0.091** | **-0.79** | **1** | **0** | **1** | **0** |
| | Thailand | 98 | 333 | 0 | 0 | NA | 0 | 0 | 1 | 0 |
| | Myanmar | 9 | 333 | 0 | 0 | NA | 0 | 0 | 1 | 0 |
| | Cambodia | 28 | 333 | 0 | 0 | NA | 0 | 0 | 1 | 0 |
| | Vietnam | 9 | 333 | 0 | 0 | NA | 0 | 0 | 1 | 0 |
| | Malaysia | 6 | 333 | 0 | 0 | NA | 0 | 0 | 1 | 0 |
| | Indonesia | 6 | 333 | 0 | 0 | NA | 0 | 0 | 1 | 0 |
| | Colombia | 30 | 333 | 0 | 0 | NA | 0 | 0 | 1 | 0 |
| | Mexico | 19 | 333 | 0 | 0 | NA | 0 | 0 | 1 | 0 |
| | Peru | 39 | 333 | 1 | 0.567 | -0.289 | 1 | 0 | 1 | 0 |

| | | | | | | | | | | |
|---|---|---|---|---|---|---|---|---|---|---|
| | Brazil | 4 | 333 | 0 | 0 | NA | 0 | 0 | 1 | 0 |
| | PNG | 14 | 333 | 0 | 0 | NA | 0 | 0 | 1 | 0 |
| *pv-fam-a* (aa 61-280) | **Global** | **264** | **1263** | **16** | **1.554** | **-0.615** | **11** | **5** | **14** | **0.608** |
| | Thailand | 98 | 1263 | 12 | 1.716 | 0.042 | 8 | 4 | 9 | 0.476 |
| | Myanmar | 9 | 1263 | 3 | 0.814 | -0.263 | 2 | 1 | 2 | 0.222 |
| | Cambodia | 29 | 1263 | 8 | 0.856 | -1.437 | 4 | 4 | 4 | 0.2 |
| | Vietnam | 10 | 1263 | 7 | 1.847 | -0.241 | 3 | 4 | 2 | 0.356 |
| | Malaysia | 6 | 1263 | 1 | 0.264 | -0.933 | 1 | 0 | 1 | 0 |
| | Indonesia | 5 | 1263 | 4 | 1.267 | -1.094 | 3 | 1 | 3 | 0.7 |
| | Colombia | 31 | 1263 | 4 | 0.96 | 0.538 | 3 | 1 | 4 | 0.63 |
| | Mexico | 19 | 1263 | 2 | 0.449 | -0.021 | 2 | 0 | 3 | 0.526 |
| | Peru | 40 | 1263 | 4 | 1.201 | 1.463 | 3 | 1 | 6 | 0.767 |
| | Brazil | 4 | 1263 | 2 | 0.924 | 0.592 | 2 | 0 | 3 | 0.833 |
| | PNG | 13 | 1263 | 5 | 1.157 | -0.331 | 3 | 2 | 3 | 0.5 |

**Table S2.** *P. vivax* reference (Sal-1) and variant (haplotype) proteins tested in immunogenicity experiments.

| PROTEIN | Region (aa) | CONSTRUCTS | | PROTEIN CONCENTRATION (mg/ml) | COUPLING AMOUNT (μg) PER $2.5x10^6$ BEADS |
|---|---|---|---|---|---|
| Pv-fam-a | 61-280 | Sal1 | PVX_096995 | 1.7 | 3 |
| | | *Variant* | SEM8 / *hap1* | 0.1728 | 8 |
| MSP8 | 24-463 | Sal1 | PVX_097625 | 0.39 | 1.4 |
| | | *Variant* | SEM10 / *hap1* | 0.1072 | 0.3 |
| RBP2b | 161-1454 | Sal1 | PVX_094255 | 1 | 0.6 |
| | | *Variant* | SEM2 / *hap1* | 0.102 | 0.3 |
| | | *Variant* | SEM4 / *hap2* | 0.126 | 0.3 |
| | | *Variant* | SEM6 / *hap3* | 0.248 | 0.7 |
| RBP2a | 160-1135 | Sal1 | PVX_121920 | 4.6 | 10 |
| | | *Variant* | SEM28 / *hap1* | 0.134 | 6 |
| | | *Variant* | SEM29 / *hap2* | 0.246 | 1.6 |
| PTEX150 | 24-908 | Sal1 | PVX_084720 | 0.24 | 2 |
| | | *Variant* | SEM39 | 0.098 | 2 |
| MSP5 | 23-365 | Sal1 | PVX_003770 | 0.58 | 0.2 |
| | | *Variant* | SEM31 / *hap1* | 0.098 | 0.2 |
| | | *Variant* | SEM33 / *hap2* | 0.093 | 0.2 |
| RIPR | 552-1075 | *Sal1* | PVX_095055 | 2.4 | 4 |
| | | *Variant* | SEM41 / *hap1* | 0.202 | 10 |
| | | *Variant* | SEM42 / *hap2* | 0.151 | 4 |
| DBPII | 242-529 | Sal1 | PVX_110810 | 1.2 | 0.6 |
| | | *AH* | AAY34130.1 | 0.6 | 1.4 |
| | | *Variant* | SEM35 / *hap1* | 0.845 | 2 |
| | | *Variant* | SEM37 / *hap2* | 0.362 | 6 |

**Table S3.** AUC (and CI) values for each *P. vivax* serological exposure marker construct tested, split via cohort. Significance was tested via bootstrap resampling.

| | | Brazil | | | | Thai | | | |
|---|---|---|---|---|---|---|---|---|---|
| Antigen_Group | **Antigen_Haplotype** | **AUC** | **CI_lower** | **CI_upper** | **P_value** | **AUC** | **CI_lower** | **CI_upper** | **P_value** |
| **DBPII** | DBPII Sal1 | 0.826939 | 0.798464 | 0.855414 | | 0.697622 | 0.593026 | 0.802217 | |
| | DBPII AH | 0.829874 | 0.801298 | 0.85845 | 0.56525 | 0.628239 | 0.515864 | 0.740615 | 0.002889 |
| | DBPII SEM-35 Hap1 | 0.816984 | 0.787805 | 0.846164 | 0.231171 | 0.723719 | 0.627227 | 0.820211 | 0.31911 |
| | DBPII SEM-37 Hap2 | 0.81614 | 0.78695 | 0.845331 | 0.200383 | 0.693058 | 0.590694 | 0.795423 | 0.870031 |
| **MSP5** | MSP5 L19 Sal1 | 0.800005 | 0.766163 | 0.833846 | | 0.821315 | 0.755189 | 0.887442 | |
| | MSP5 SEM-31 | 0.811776 | 0.778393 | 0.84516 | 0.624128 | 0.802511 | 0.720279 | 0.884744 | 0.726553 |
| | MSP5 SEM-33 | 0.81673 | 0.783895 | 0.849565 | 0.477528 | 0.770219 | 0.672888 | 0.867549 | 0.392538 |
| **MSP8** | MSP8 L34 Sal1 | 0.839217 | 0.811072 | 0.867361 | | 0.783425 | 0.690458 | 0.876393 | |
| | MSP8 SEM-10 Hap1 | 0.812562 | 0.781776 | 0.843347 | 4E-05 | 0.685914 | 0.575798 | 0.796029 | 8.2E-08 |
| **RBP2A** | RBP2a Sal1 | 0.847987 | 0.819884 | 0.876091 | | 0.70475 | 0.601057 | 0.808442 | |
| | RBP2a SEM-28 Hap1 | 0.804561 | 0.773742 | 0.83538 | 7.84E-05 | 0.787473 | 0.695249 | 0.879696 | 0.00197 |
| | RBP2a SEM-29 Hap9 | 0.764375 | 0.727811 | 0.800939 | 3.19E-09 | 0.69116 | 0.579753 | 0.802567 | 0.652429 |
| **RBP2B** | RBP2b P25 | 0.829181 | 0.793897 | 0.864465 | | 0.833517 | 0.768002 | 0.899032 | |
| | RBP2b SEM-2 | 0.84352 | 0.812 | 0.875039 | 0.128115 | 0.822733 | 0.756485 | 0.888982 | 0.150054 |
| | RBP2b SEM-4 | 0.843536 | 0.810927 | 0.876144 | 0.07035 | 0.82115 | 0.752395 | 0.889904 | 0.143807 |
| | RBP2b SEM-6 | 0.849574 | 0.818713 | 0.880435 | 0.038641 | 0.829331 | 0.765088 | 0.893575 | 0.63428 |
| **RIPR** | RIPR | 0.861602 | 0.83575 | 0.887454 | | 0.734463 | 0.641976 | 0.82695 | |
| | RIPR SEM-41 | 0.832971 | 0.803036 | 0.862906 | 0.16162 | 0.636583 | 0.516448 | 0.756718 | 0.188805 |
| | RIPR SEM-42 | 0.829998 | 0.79981 | 0.860185 | 0.122901 | 0.649657 | 0.533465 | 0.765849 | 0.250339 |
| **PTEX150** | PTEX150 L18 Sal1 | 0.765437 | 0.731339 | 0.799535 | | 0.733096 | 0.629611 | 0.83658 | |
| | PTEX150 SEM-39 Hap1 | 0.762944 | 0.729106 | 0.796781 | 0.571801 | 0.751349 | 0.648136 | 0.854562 | 0.232627 |
| **Fam-a** | Pv-fam-a L02 Sal1 | 0.816314 | 0.786588 | 0.84604 | | 0.739941 | 0.642694 | 0.837188 | |
| | Pv-fam-a SEM-8 Hap1 | 0.812753 | 0.782412 | 0.843094 | 0.419635 | 0.756179 | 0.66413 | 0.848228 | 0.290575 |

**Table S4.** Top antigen combinations (ranging from two to eight) for the combined dataset (Brazil, Thailand and Negative Controls), as well as the Brazil and Thailand cohort datasets. All selected antigen combinations include a haplotype of RBP2a (RBP2a.sal1, SEM.28 or SEM.29). Classification performance of the random forest model for each combination is presented as the area under the curve (AUC) along with the standard error.

| Top | Combination (*Bold are haplotypes for RBP2a) | AUC | Standard Error |
|---|---|---|---|
| Combined Dataset | | | |
| 2 | **RBP2a.sal1** + MSP8 | 0.843 | 0.007 |
| 3 | **RBP2a .sal1**+ RBP2b + MSP8 | 0.870 | 0.007 |
| 4 | **RBP2a.sal1** + RBP2b + PvRIPR + MSP8 | 0.878 | 0.006 |
| 5 | **RBP2a.sal1** + RBP2b + PvRIPR + PV-FAM-A + MSP8 | 0.882 | 0.006 |
| 6 | **RBP2a.sal1** + MSP5 + RBP2b + PvRIPR + PV-FAM-A + MSP8 | 0.886 | 0.006 |
| 7 | **RBP2a.sal1** +MSP5 + RBP2b + Pv.DBPII.sal1 + PvRIPR + PV-FAM-A + MSP8 | 0.885 | 0.007 |
| 8 | **RBP2a.sal1** + MSP5 + RBP2b + Pv.DBPII.sal1 + PvRIPR + PTEX150 + PV-FAM-A + MSP8 | 0.881 | 0.007 |
| 2 | **SEM.28** + RBP2b | 0.820 | 0.006 |
| 3 | **SEM.28** + RBP2b + MSP8 | 0.867 | 0.006 |
| 4 | **SEM.28** + MSP5 + RBP2b + MSP8 | 0.873 | 0.006 |
| 5 | **SEM.28** + RBP2b + PvRIPR + PV-FAM-A + MSP8 | 0.881 | 0.006 |
| 6 | **SEM.28** + MSP5 + RBP2b + PvRIPR + PV-FAM-A + MSP8 | 0.886 | 0.006 |
| 7 | **SEM.28** + MSP5 + RBP2b + Pv.DBPII.sal1 + PvRIPR + PV-FAM-A + MSP8 | 0.884 | 0.006 |
| 8 | **SEM.28** +MSP5 + RBP2b + Pv.DBPII.sal1 + PvRIPR + PTEX150 + PV-FAM-A + MSP8 | 0.879 | 0.007 |
| 2 | **SEM.29** + PvRIPR | 0.796 | 0.007 |
| 3 | **SEM.29** + RBP2b + MSP8 | 0.861 | 0.006 |
| 4 | **SEM.29** + RBP2b + Pv.DBPII.sal1 + MSP8 | 0.877 | 0.006 |
| 5 | **SEM.29** + MSP5 + RBP2b + Pv.DBPII.sal1 + MSP8 | 0.878 | 0.007 |
| 6 | **SEM.29** + PvRIPR + MSP5 + RBP2b + PV-FAM-A + MSP8 | 0.883 | 0.006 |
| 7 | **SEM.29** + MSP5 + RBP2b + Pv.DBPII.sal1 + PvRIPR + PV-FAM-A + MSP8 | 0.882 | 0.006 |
| 8 | **SEM.29** + MSP5 + RBP2b + Pv.DBPII.sal1 + PvRIPR + PTEX150 + PV-FAM-A + MSP8 | 0.878 | 0.007 |
| Brazil Dataset | | | |
| 2 | **RBP2a.sal1** + MSP8 | 0.795 | 0.008 |
| 3 | **RBP2a.sal1** + RBP2b + MSP8 | 0.835 | 0.007 |
| 4 | **RBP2a.sal1** + RBP2b + PV-FAM-A + MSP8 | 0.847 | 0.007 |
| 5 | **RBP2a.sal1** + RBP2b + Pv.DBPII.sal1 + PV-FAM-A + MSP8 | 0.855 | 0.007 |
| 6 | **RBP2a.sal1** + RBP2b + Pv.DBPII.sal1 + PvRIPR + PV-FAM-A + MSP8 | 0.858 | 0.006 |
| 7 | **RBP2a.sal1** +MSP5 + RBP2b + Pv.DBPII.sal1 + PvRIPR + PV-FAM-A + MSP8 | 0.858 | 0.006 |
| 8 | **RBP2a.sal1** + MSP5 + RBP2b + Pv.DBPII.sal1 + PvRIPR + PTEX150 + PV-FAM-A + MSP8 | 0.856 | 0.006 |
| 2 | **SEM.28** + PvRIPR | 0.786 | 0.006 |
| 3 | **SEM.28** + RBP2b + MSP8 | 0.841 | 0.007 |
| 4 | **SEM.28** + RBP2b + PV-FAM-A + MSP8 | 0.850 | 0.007 |
| 5 | **SEM.28** + RBP2b + PvRIPR + PV-FAM-A + MSP8 | 0.856 | 0.006 |
| 6 | **SEM.28** + MSP5 + RBP2b + PvRIPR + PV-FAM-A + MSP8 | 0.858 | 0.006 |
| 7 | **SEM.28** + MSP5 + RBP2b + Pv.DBPII.sal1 + PvRIPR + PV-FAM-A + MSP8 | 0.857 | 0.006 |
| 8 | **SEM.28** + MSP5 + RBP2b + Pv.DBPII.sal1 + PvRIPR + PTEX150 + PV-FAM-A + MSP8 | 0.852 | 0.007 |
| 2 | **SEM.29** + RBP2b | 0.774 | 0.007 |
| 3 | **SEM.29** + RBP2b + MSP8 | 0.836 | 0.007 |

| | | | |
|---|---|---|---|
| 4 | **SEM.29** + RBP2b + PV-FAM-A + MSP8 | 0.848 | 0.007 |
| 5 | **SEM.29** + RBP2b + PvRIPR + PV-FAM-A + MSP8 | 0.856 | 0.007 |
| 6 | **SEM.29** + MSP5 + RBP2b + PvRIPR + PV-FAM-A + MSP8 | 0.859 | 0.006 |
| 7 | **SEM.29** + MSP5 + RBP2b + Pv.DBPII.sal1 + PvRIPR + PV-FAM-A + MSP8 | 0.859 | 0.006 |
| 8 | **SEM.29** + MSP5 + RBP2b + Pv.DBPII.sal1 + PvRIPR + PTEX150 + PV-FAM-A + MSP8 | 0.855 | 0.006 |
| Thailand Dataset | | | |
| 2 | **RBP2a.sal1** + RBP2b | 0.752 | 0.024 |
| 3 | **RBP2a.sal1** + RBP2b + MSP8 | 0.807 | 0.026 |
| 4 | **RBP2a.sal1** + RBP2b + PV-FAM-A + MSP8 | 0.823 | 0.022 |
| 5 | **RBP2a.sal1** + RBP2b + PvRIPR + PV-FAM-A + MSP8 | 0.829 | 0.023 |
| 6 | **RBP2a.sal1** + RBP2b + Pv.DBPII.sal1 + PvRIPR + PV-FAM-A + MSP8 | 0.817 | 0.028 |
| 7 | **RBP2a.sal1** + RBP2b + Pv.DBPII.sal1 + PvRIPR + PTEX150 + PV-FAM-A + MSP8 | 0.797 | 0.028 |
| 8 | **RBP2a.sal1** + MSP5 + RBP2b + Pv.DBPII.sal1 + PvRIPR + PTEX150 + PV-FAM-A + MSP8 | 0.780 | 0.029 |
| 2 | **SEM.28** + RBP2b | 0.759 | 0.026 |
| 3 | **SEM.28** + RBP2b + PV-FAM-A | 0.787 | 0.023 |
| 4 | **SEM.28** + RBP2b + PvRIPR + MSP8 | 0.820 | 0.023 |
| 5 | **SEM.28** + RBP2b + PvRIPR + PV-FAM-A + MSP8 | 0.836 | 0.023 |
| 6 | **SEM.28** + MSP5 + RBP2b + PvRIPR + PV-FAM-A + MSP8 | 0.822 | 0.025 |
| 7 | **SEM.28** + MSP5 + RBP2b + Pv.DBPII.sal1 + PvRIPR + PV-FAM-A + MSP8 | 0.802 | 0.027 |
| 8 | **SEM.28** + MSP5 + RBP2b + Pv.DBPII.sal1 + PvRIPR + PTEX150 + PV-FAM-A + MSP8 | 0.783 | 0.028 |
| 2 | **SEM.29** + RBP2b | 0.787 | 0.023 |
| 3 | **SEM.29** + RBP2b + MSP8 | 0.820 | 0.022 |
| 4 | **SEM.29** + RBP2b + PvRIPR + MSP8 | 0.822 | 0.022 |
| 5 | **SEM.29** + RBP2b + PvRIPR + PV-FAM-A + MSP8 | 0.827 | 0.024 |
| 6 | **SEM.29** + RBP2b + PvRIPR + PTEX150 + PV-FAM-A + MSP8 | 0.808 | 0.022 |
| 7 | **SEM.29** + RBP2b + Pv.DBPII.sal1 + PvRIPR + PTEX150 + PV-FAM-A + MSP8 | 0.801 | 0.024 |
| 8 | **SEM.29** + MSP5 + RBP2b + Pv.DBPII.sal1 + PvRIPR + PTEX150 + PV-FAM-A + MSP8 | 0.778 | 0.028 |
